# Local control of dopamine release in nucleus accumbens gates opioid withdrawal aversion

**DOI:** 10.64898/2026.05.25.727738

**Authors:** Matthew B. Pomrenze, Jason. M. Tucciarone, Gavin C. Touponse, Nicholas Denomme, BaDoi N. Phan, Robyn St. Laurent, Julia A. Galiza Soares, Daniel F. Cardozo Pinto, Michaela Y. Guo, Jinhee Baek, Allen P.F. Chen, Zihui Zhang, Amei Shank, Zachary Freyberg, Andreas R. Pfenning, Neir Eshel, Robert C. Malenka

## Abstract

The opioid crisis has emphasized the need for more effective treatments for opioid use disorder (OUD)^1–3^, which is characterized by habitual opioid use to avoid withdrawal symptoms^4,5^. Both physical and affective symptoms contribute to opioid withdrawal yet whether different neural mechanisms mediate these different symptom domains and contribute distinctly to opioid relapse is unknown. While neurons expressing mu opioid receptors (MORs) gate opioids’ reinforcing effects^6–8^ by increasing dopamine (DA) release in nucleus accumbens (NAc), sharp decreases in NAc DA release are associated with withdrawal^9–11^, the cellular and circuit mechanisms of which are unknown. Here we describe an unusual population of evolutionarily-conserved MOR+ neurons in the NAc expressing the transcription factor *Tshz1*. Increased activity in these neurons is required for withdrawal aversion learning. Deletion of MORs in *Tshz1* neurons prevented withdrawal-induced decreases in DA release and affective aversion, but not physical symptoms associated with withdrawal. Pharmacological activation of mGluR8, which is preferentially expressed in *Tshz1* neurons, reduced withdrawal aversion. Thus, by dissociating the circuit mechanisms contributing to the physical and affective components of opioid withdrawal focusing on the critical role of *Tshz1* neurons, we have identified a novel druggable target with therapeutic potential for treating key OUD withdrawal symptoms.

## Main text

The opioid crisis has surged over the past two decades, leading to a critical need for innovative treatments for opioid use disorder (OUD)^1–3^. The pathogenesis of OUD is characterized by devastating cycles of opioid intoxication and withdrawal which lead to dependence. Withdrawal presents as a spectrum of physical (e.g. fever, nausea, pain, escape behaviors) and affective (e.g. aversion, dysphoria, agitation, craving) symptoms, which together promote habitual opioid use^4,5^, thereby perpetuating the cycles of intoxication and withdrawal characteristic of OUD. It remains unclear whether the physical and affective symptoms elicited by withdrawal are mediated by distinct neural mechanisms, which play unique roles in the progression of OUD.

The effects of opioids are largely mediated through the cells in which MORs are expressed^6–8^. Dopamine (DA) release in the NAc, via MOR-induced disinhibition of ventral tegmental area (VTA) DA neurons, mediates the rewarding properties of opioids^7,12–15^, while suppression of DA release appears to accompany withdrawal^10,11,16^. Although direct inhibition of NAc DA release is commonly observed to be aversive^17–19^, the mechanisms mediating this effect on withdrawal are unclear as is its relationship to physical withdrawal symptoms, which involve several other brain regions including the central amygdala^9^, thalamus^20^, prefrontal cortex^21^ and locus coeruleus^22–24^.

Here we focus on the role of the NAc in opioid withdrawal because it is enriched in MORs^25,26^, has cells that are activated during opioid withdrawal^27,28^, and direct infusion of opioid antagonists into the NAc produces withdrawal aversion^29^. Using a combination of behavioral, genetic, and physiological approaches, we discover a previously unknown and evolutionary conserved cell-type in the NAc that is crucial for the aversive nature of opioid withdrawal through modulation of local NAc DA release, but not withdrawal-elicited physical symptoms. Single-cell gene expression analyses revealed a novel molecular target in these NAc *Tshz1* cells, the pharmacological engagement of which robustly blocked withdrawal aversion.

### Opioid withdrawal triggers robust suppression of DA release in the NAc

Early dialysis studies detected reductions in DA release in the NAc medial shell during acute opioid withdrawal^10,11^. To test the replicability of this result, we expressed the DA sensor GRAB DA in the NAc medial shell of wild-type mice and performed fiber photometry recordings while precipitating opioid withdrawal in mice that were made opioid dependent over 5 days^12^ (Fig. 1a,b). On day 6, mice received a final dose of morphine (or saline) during a photometry recording and 40 min later were injected with naloxone (Fig. 1b). Morphine elicited a surge of DA, followed by a robust suppression of release below baseline in response to naloxone while saline generated no detectable changes (Fig. 1c-e and Extended Data Fig. 1a-c). Mice administered the same morphine-naloxone dose regimen in a conditioned place aversion (CPA) assay learned to avoid the context associated with withdrawal, indicating a profoundly aversive experience (Fig. 1f-h). Thus, the naloxone-elicited suppression of DA release is associated with the naloxone-mediated aversive nature of withdrawal.

**Fig 1.**
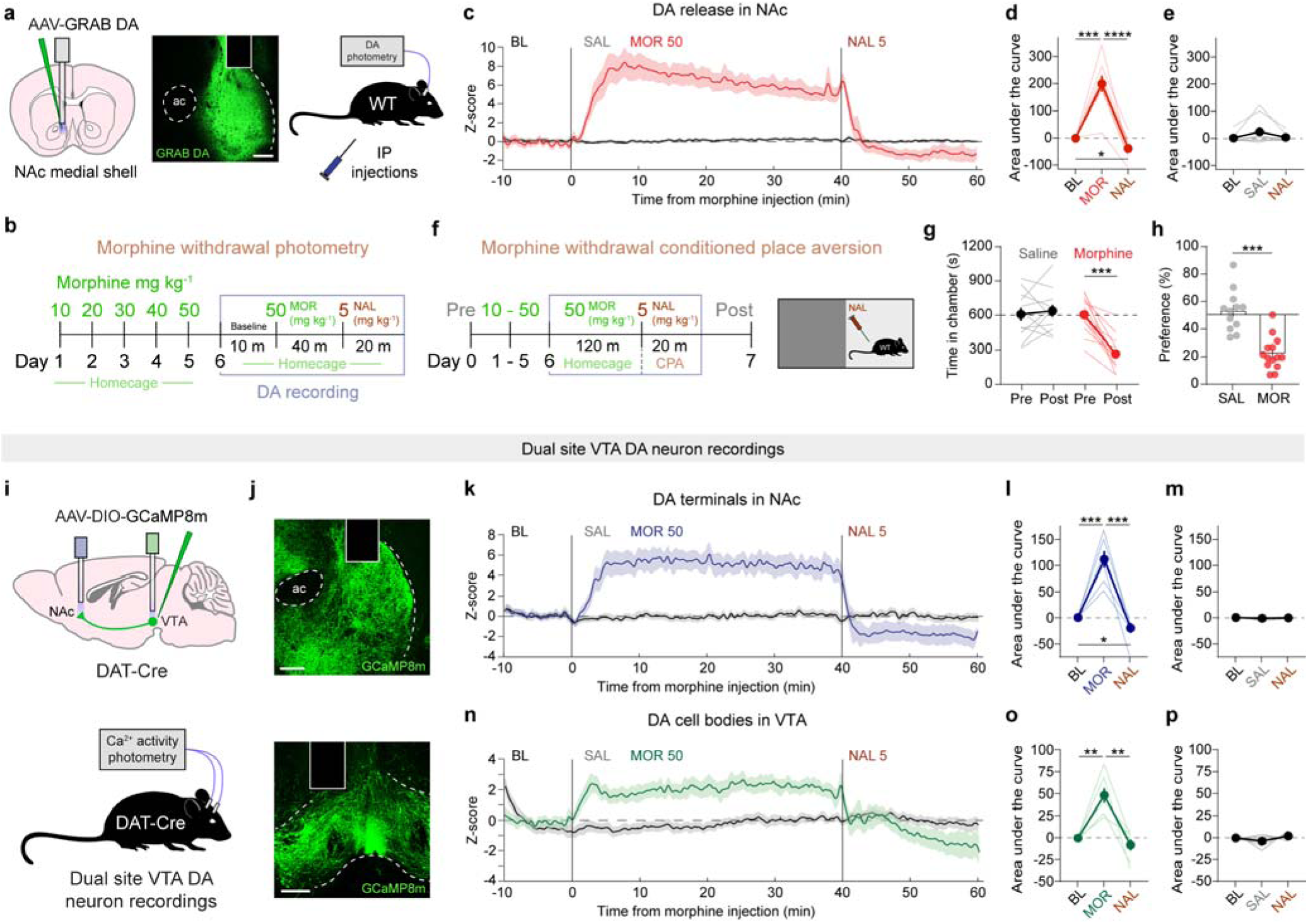
Opioid withdrawal triggers robust suppression of DA release in the NAc. **a.** Schematic and image of GRAB DA recording during precipitated opioid withdrawal. Scale = 100 um. **b.** Time course of experimental design. Mice were made morphine dependent in the home cage and then put into withdrawal during a photometry recording. Green denotes days/times with morphine administration and red with naloxone. **c.** Bulk DA release in the NAc during morphine (50 mg kg^-1^) or saline administration and precipitated withdrawal with naloxone (5 mg kg^-1^). **d.** Area under the curve for each drug phase. RM one-way ANOVA, F_2,24_ = 63.25, ****p < 0.0001. Dunnett’s multiple comparisons test, BL vs MOR ***p = 0.0002, MOR vs NAL, ****p < 0.0001, BL vs NAL *p = 0.0388. *n* = 9 mice. **e.** Area under the curve for each drug phase after saline injection. RM one-way ANOVA, F_2,24_ = 2.843, p = 0.1285. *n* = 9 mice. **f.** Timeline and schematic of morphine withdrawal conditioned place aversion (CPA) assay. **g.** Compared with mice administered saline, morphine-dependent mice avoid the chamber paired with naloxone. RM two-way ANOVA, time x drug F_1,24_ = 25.15, ****p < 0.0001. Sidak’s multiple comparisons test, Saline: Pre vs Post, p = 0.8264, Morphine: Pre vs Post, ****p < 0.0001. Saline *n* = 12 mice, morphine *n* = 14 mice. **h.** Direct comparison of CPA performance between saline and morphine groups. Unpaired, two-tailed t-test, t_24_ = 5.892, ****p < 0.0001. **i.** Top, schematic of dual calcium recordings in NAc and VTA. Bottom, schematic of DAT-Cre mouse with the dual site VTA DA neuron recording. **j.** Top, image of recording site with DA terminals in the NAc. Bottom, image of recording site with DA cell bodies in the NAc from the same mouse. Scale = 100 um. **k.** Average bulk calcium activity in DA axon terminals in the NAc during morphine (50 mg kg^-1^) administration and precipitated withdrawal with naloxone (5 mg kg^-1^). **l.** Area under the curve of each drug phase for terminal recordings. RM one-way ANOVA, F_2,18_ = 47.38, ****p < 0.0001. Dunnett’s multiple comparisons test, BL vs MOR ***p = 0.0007, MOR vs NAL, ***p < 0.0005, BL vs NAL p = 0.142. BL vs NAL, paired, two-tailed wilcoxon matched-paired signed rank test, *p = 0.0156. *n* = 7 mice. **m.** Area under the curve after saline injection. RM one-way ANOVA, F_2,18_ = 0.4597, p = 0.5971. *n* = 7 mice. **n.** Average bulk calcium activity in DA cell bodies in the VTA during morphine (50 mg kg^-1^) administration and precipitated withdrawal with naloxone (5 mg kg^-1^). **o.** Area under the curve of each drug phase for cell body recordings. RM one-way ANOVA, F_2,18_ = 25.72, ****p = 0.0005. Dunnett’s multiple comparisons test, BL vs MOR **p = 0.0017, MOR vs NAL, **p = 0.0036, BL vs NAL p = 0.373. BL vs NAL paired, two-tailed wilcoxon matched-paired signed rank test, p = 0.6875. *n* = 7 mice. **p.** Area under the curve after saline injection. RM one-way ANOVA, F_2,18_ = 0.1172, p = 0.3305. *n* = 7 mice.

To determine the mechanism underlying this inhibition of DA release, we performed simultaneous recordings of VTA DA neuron cell bodies and axon terminals in the NAc by injecting AAV-DIO-GCaMP8m into VTA of DAT-Cre mice and implanting optical fibers over both the VTA and ipsilateral NAc medial shell (Fig. 1i,j). Simultaneous recordings revealed increased activity in both DA axon terminals (Fig. 1k-m and Extended Data Fig. 1d,e) and DA cell bodies (Fig. 1n-p and Extended Data Fig. 1d,f) after morphine injection. Cross-correlation analysis identified a significant increase in VTA DA cell body and axon synchronization in response to morphine, suggesting cell bodies as the source of activity in the axons (Extended Data Fig. 1g,h,j,k). Naloxone suppressed activity in both cellular compartments (Fig. 1k-p). Terminal activity rapidly shut down and was completely silent within 2-3 minutes, with DA transients slowly reemerging about 10 min later (possibly reflecting the short half-life of naloxone), although the bulk signal remained beneath baseline levels (Fig. 1k and Extended Data Fig. 1e). Cell body activity decreased at a slower rate (Fig. 1n and Extended Data Fig. 1f). Cross-correlation analysis of the naloxone phase showed a reduction in the peak correlation compared with morphine and baseline phases, indicating decoupling of DA cell bodies and axons (Extended Data Fig. 1i-k). These data show that DA release is robustly inhibited during acute opioid withdrawal, possibly through multiple inhibitory sources modulating DA cell body and terminal activity.

### Midbrain GABA neurons promote opioid reward, but not withdrawal

The acute rewarding effect of opioids is thought to be due to disinhibition of VTA DA neuron activity via MOR-mediated inhibition of GABA neurons in the caudal VTA and rostral medial tegmental nucleus (RMTg)^9,12–14,30^. Consistent with this hypothesis, morphine decreased GCaMP8m activity in GABA neurons in the caudal midbrain (Extended Data Fig. 2a,b) and naloxone administration returned GABA neuron activity to baseline levels (Extended Data Fig. 2c,d). No effect of naloxone on GABA neuron activity was observed after saline injections (Extended Data Fig. 2c,e). To determine if activity changes in these GABA cells contributed to both opioid reward and withdrawal aversion, we genetically deleted *Oprm1* (gene encoding MORs) in the caudal midbrain by injecting AAV-Cre-eGFP into this region of floxed *Oprm1* (*Oprm1*^fl/fl^) mice and tested for opioid reward in a morphine-elicited conditioned place preference (CPP) procedure (Extended Data Fig. 2f,g). Deletion of *Oprm1* in the midbrain robustly disrupted morphine CPP (Extended Data Fig. 2h) and locomotor sensitization to morphine (Extended Data Fig. 2i). In contrast, deletion of *Oprm1* had no effect on avoidance of the withdrawal-paired chamber in the CPA assay (Extended Data Fig. 2j-l). To determine whether physical withdrawal symptoms were affected by *Oprm1* deletion, we prepared a new cohort and monitored jumping, rearing, and tremors during withdrawal (Extended Data Fig. 2m). Again, no differences in the expression of these physical signs were detected (Extended Data Fig. 2n). These data suggest that inhibition of midbrain MOR GABA neurons mediate the acute reinforcing effects of opioid reward, but that their responses to opioids during repeated administration are not required for development of either the physical symptoms of withdrawal nor the aversive nature of withdrawal.

### Identification of *Oprm1*-enriched NAc *Tshz1* neurons

We next focused on the potential role of NAc neurons in mediating withdrawal symptoms because they exhibit cFos induction during withdrawal^27–29^ as well as robust expression of *Oprm1*^25,26^. Prior single-nuclei RNA sequencing (snRNAseq) of NAc from naïve mice^31,32^ (Fig. 2a) revealed three distinct clusters of neurons (Fig. 2b) based on the neuronal “archetypes” found in the striatum^33^.

**Fig 2.**
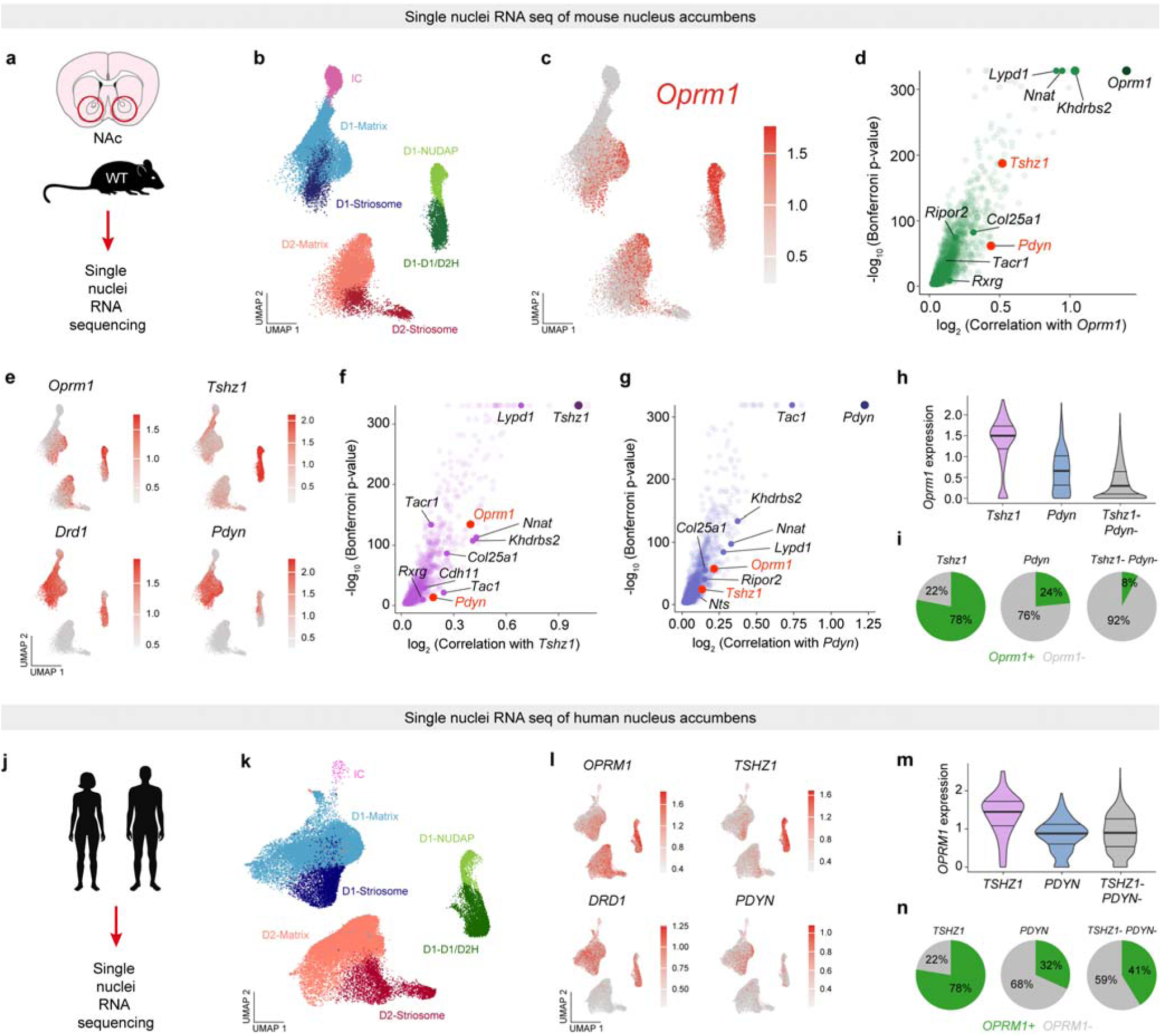
NAc *Tshz1* neurons are enriched with *Oprm1* in mice and humans. **a.** Single nuclei RNAseq of mouse NAc. **b.** Uniform manifold approximation and projection (UMAP) dimensional reduction and visualization of transcriptional profiles of NAc neurons. Colors represent sub-clustering based on previous characterization of striatal cell “archetypes”. **c.** Feature plot showing the expression pattern of *Oprm1*. **d.** Volcano plot showing genes significantly correlated with *Oprm1* expression. **e.** Feature plots showing the distributions of other key marker genes in NAc. **f.** Volcano plot showing genes significantly correlated with *Tshz1* expression. **g.** Volcano plot showing genes significantly correlated with *Pdyn* expression. **h.** Violin plot quantifying the expression of *Oprm1* in each cell-type. Mixed effect linear model, *Tshz1* (NUDAP) vs *Pdyn* ****p = 9.465e-53, *Tshz1* (NUDAP) vs *Tshz1*- *Pdyn*- ****p = 5.1551e-228. *Tshz1* (NUDAP) vs *Pdyn,* Bonferroni corrected p = 1.8932e-52. *Tshz1* (NUDAP) vs *Tshz1*- *Pdyn*-, Bonferroni corrected p = 1.031e-227. Horizontal lines indicate 25%, 50%, and 75% quartiles. **i.** Pie charts illustrating the proportion of each cell-type expressing *Oprm1*. **j.** Single nuclei RNAseq of human NAc. **k.** UMAP dimensional reduction and visualization of transcriptional profiles of human NAc neurons. Colors represent sub-clustering of striatal “archetypes”, similar to mouse NAc. **l.** Feature plots showing the distributions of key marker genes in human NAc. **m.** Violin plot quantifying the expression of *OPRM1* in each cell-type. Mixed effect linear model, *TSHZ1* (NUDAP) vs *PDYN* ****p = 1.0634e-52, *TSHZ1* (NUDAP) vs *TSHZ1*- *PDYN*- ****p = 5.0318e-67. *TSHZ1* (NUDAP) vs *PDYN,* Bonferroni corrected p = 2.1268e-52. *TSHZ1* (NUDAP) vs *TSHZ1*- *PDYN*-, Bonferroni corrected p = 1.0063e-75. Horizontal lines indicate 25%, 50%, and 75% quartiles. **n.** Pie charts illustrating the proportion of each cell-type expressing *OPRM1*.

Subclusters included cells belonging to D1 or D2 dopamine receptor-expressing populations, striosome- or matrix-like populations, eccentric D1 cells with a D1/D2 hybrid subpopulation and one previously termed “neurochemically unique domains in the accumbens and putamen” (NUDAP)^33^. There was a wide distribution of *Oprm1* expression, with the highest expression and density in the eccentric D1 cluster (Fig. 2c). A search for genes whose expression significantly correlated with *Oprm1* revealed genes previously found in cells located in striosomes^34^, including *Lypd1*, *Nnat*, *Tshz1*, and *Pdyn* (Fig. 2d).

We focused on NAc neurons expressing *Tshz1* (Teashirt zinc finger homeobox 1^35^), as *Tshz1* neurons in the dorsal striatum are important for aversion learning^36^. Consistent with our analysis, recent single-cell RNA sequencing studies have identified *Tshz1* as a marker for neurons in striosomes^37^ and neurons enriched with *Oprm1* in the rat NAc^25^. We mapped *Tshz1* expression onto the NAc cell clusters and found the greatest expression and density in the eccentric D1 cluster (Fig. 2e). We also mapped *Drd1* and *Pdyn* (a classic D1 marker gene encoding dynorphin) and found *Drd1* highly expressed in both the large D1 cluster and the eccentric cluster, while *Pdyn* primarily localized only to the D1 cluster and preferentially in D1-striosome (Fig. 2e). As expected, *Drd2* and *Penk* localized to the large D2 cluster (Extended Data Fig. 3b). The highest degree of *Oprm1* and *Tshz1* co-expression was observed in the D1-NUDAP archetype, associated with low levels of *Pdyn* (Extended Data Fig. 3c-g). Furthermore, *Oprm1* expression was significantly correlated with *Tshz1* expression (Fig. 2f), but much less so with *Pdyn* (Fig. 2f,g). Thus, NUDAP *Tshz1* neurons were significantly more enriched with *Oprm1* than *Pdyn* neurons (Fig. 2h,i) and *Tshz1* neurons of the NUDAP archetype appear to be the most *Oprm1*-enriched cell-types in the NAc.

To determine whether this *Oprm1*-enriched *Tshz1* cell-type is evolutionarily conserved, we analyzed snRNAseq data from human NAc tissue^31^ (Fig. 2j). This analysis revealed a similar distribution of cell-types (Fig. 2k), with *TSHZ1* neurons significantly more enriched with *OPRM1* than *PDYN* neurons or those without either marker (Fig. 2l,m). Human NAc cells co-expressing large amounts of *TSHZ1* and *OPRM1* mapped onto the D1/D2 hybrid and NUDAP archetypes (Extended Data Fig. 3j,m), while cells enriched with *PDYN* mapped onto D1-striosome and D1-matrix archetypes and expressed low levels of *TSHZ1* and *OPRM1* (Extended Data Fig. 3k-m). The enrichment of *OPRM1* in human *TSHZ1* neurons was similar to mouse, with a similar number of *TSHZ1* cells expressing higher levels of *OPRM1* than *PDYN* cells or any other cell-type (Fig. 2m,n; Extended Data Fig. 3m,n). Overall, the *OPRM1*-enriched *TSHZ1* cell-type found in human NAc was strikingly similar to mouse. These transcriptomic data indicate *Tshz1* neurons as an evolutionary-conserved, opioid-sensitive cell-type.

### NAc *Tshz1* neurons mediate withdrawal aversion

We next determined the spatial location of *Oprm1*-enriched *Tshz1* neurons using fluorescence *in situ* hybridization of *Oprm1*, *Tshz1*, and *Pdyn* across NAc subregions (Fig. 3a and Extended Data Fig. 4a-d). *Tshz1* neurons in the medial shell were the most enriched with *Oprm1* (Fig. 3b-d) with *Pdyn* neurons expressing significantly less *Oprm1* (Fig. 3d). Consistent with our snRNAseq data, *Tshz1* and *Pdyn* cell-types were mostly distinct from one another (Extended Data Fig. 4e). However, we identified strong co-localization of *Oprm1*, *Tshz1,* and *Pdyn* in the dorsal striatum forming striosome clusters and in the lateral edge of the ventral striatum forming the “lateral strip” (Extended Data Fig. 4f,g). These expression patterns suggest a unique functional role for *Tshz1* neurons in the NAc medial shell.

**Fig 3.**
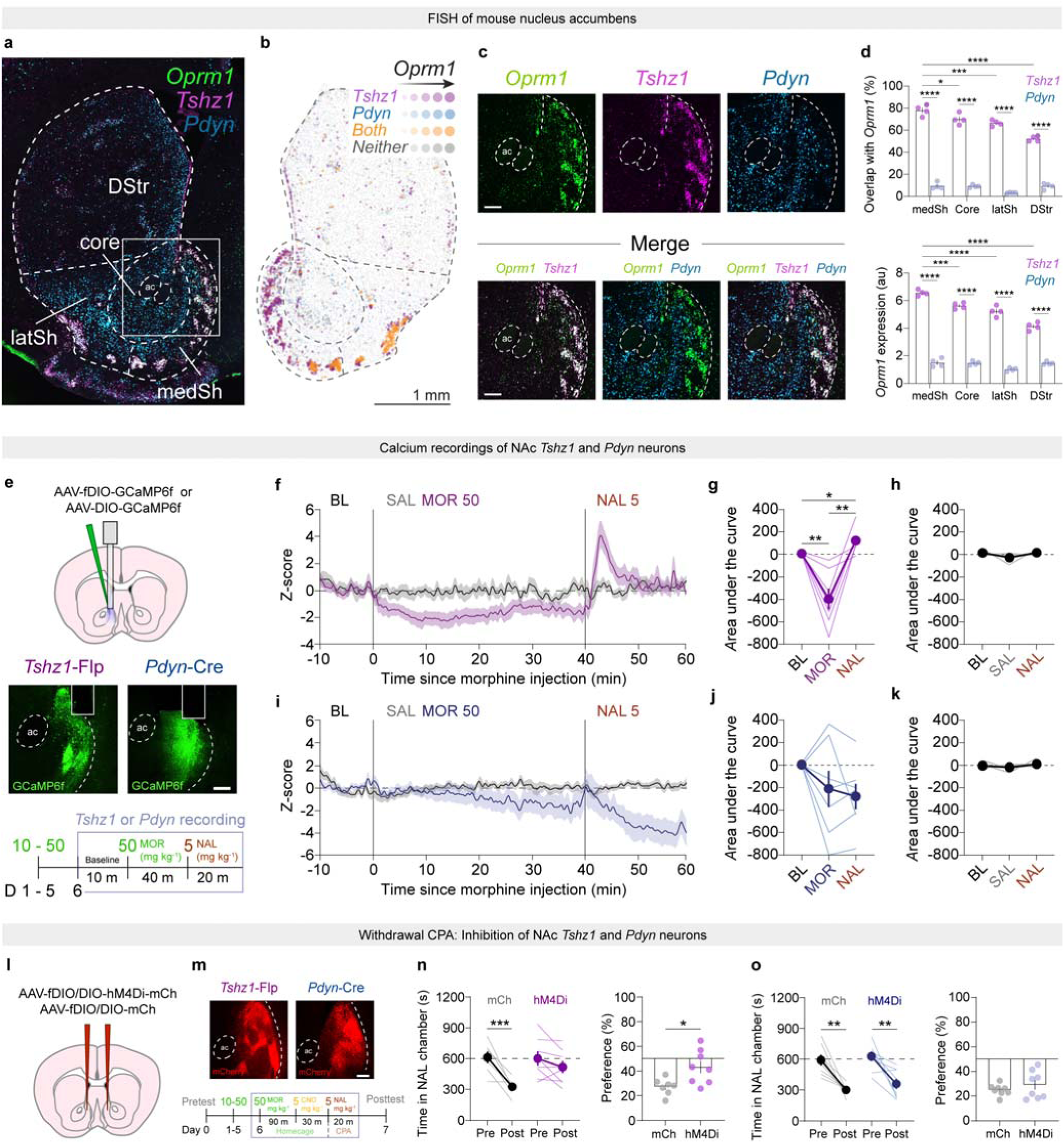
NAc *Tshz1* neuron activity mediates opioid withdrawal aversion. **a.** Fluorescence *in situ* hybridization of *Oprm1*, *Tshz1*, and *Pdyn* in mouse striatum. **b.** Reconstruction of cells across the NAc and dorsal striatum, scaled based on *Oprm1* expression. Scale = 1 mm. **c.** Region of interest in panel a. Top, individual fluorescence channels. Bottom, channel merges comparing *Oprm1* expression with the two cell-types. Scales = 100 um. **d.** Top, quantification of overlap (co-localization) with *Oprm1*. Two-way ANOVA, gene x subregion F_3,24_ = 24.89, ****p < 0.0001. Tukey’s multiple comparisons, *Tshz1* vs *Pdyn*, medSh ****p < 0.0001, Core ****p < 0.0001, latSh ****p < 0.0001, DStr ****p < 0.0001. *Tshz1* medSh vs Core *p = 0.026, *Tshz1* medSh vs latSh ***p = 0.0006, *Tshz1* medSh vs DStr ****p < 0.0001. Bottom, quantification of *Oprm1* expression in NAc *Tshz1* and *Pdyn* neurons. Two-way ANOVA, gene x subregion F_3,24_ = 35.07, ****p < 0.0001. Tukey’s multiple comparisons, *Tshz1* vs *Pdyn*, medSh ****p < 0.0001, Core ***p = 0.0002, latSh ****p < 0.0001, DStr ****p < 0.0001. *Tshz1* medSh vs Core *p = 0.026, *Tshz1* medSh vs latSh ***p = 0.0006, *Tshz1* medSh vs DStr ****p < 0.0001. *n* = 4 wild-type mice. **e.** Top, schematic of recordings of *Tshz1* and *Pdyn* neurons in NAc. Center, representative images of GCaMP6f expression in NAc medial shell *Tshz1* or *Pdyn* neurons, scale = 100 um. Bottom, time course of photometry recordings during precipitated withdrawal. **f.** Average bulk activity of NAc *Tshz1* neurons during morphine (50 mg kg^-1^) or saline administration and precipitated withdrawal with naloxone (5 mg kg^-1^). **g.** Area under the curve of each drug phase for *Tshz1* neuron recordings. RM one-way ANOVA, F_2,18_ = 15.44, **p = 0.0027. Dunnett’s multiple comparisons test, BL vs MOR *p = 0.0204, MOR vs NAL, *p = 0.0168, BL vs NAL p = 0.9704. *n* = 7 mice. **h.** Area under the curve for each drug phase after injection of saline. RM one-way ANOVA, F_2,9_ = 1.609, p = 0.2935. *n* = 4 mice. **i.** Average bulk activity of NAc *Pdyn* neurons during morphine (50 mg kg^-1^) or saline administration and precipitated withdrawal with naloxone (5 mg kg^-1^). **j.** Area under the curve of each drug phase for *Pdyn* neuron recordings. RM one-way ANOVA, F_2,18_ = 2.524, p = 0.1376. Dunnett’s multiple comparisons test, BL vs MOR p = 0.3686, MOR vs NAL, p = 0.7867, BL vs NAL p = 0.0756. *n* = 7 mice. **k.** Area under the curve for each drug phase after injection of saline. RM one-way ANOVA, F_2,12_ = 0.791, p = 0.4644. *n* = 5 mice. **l.** Schematic of AAV injections into the NAc of *Tshz1*-Flp and *Pdyn*-Cre mice. **m.** Top, images of hM4Di-mCherry expression in NAc medial shell *Tshz1* or *Pdyn* neurons, scale = 100 um. Bottom, CPA time course and experimental design. **n.** Left, mice with inhibition of *Tshz1* neurons fail to avoid the chamber paired with naloxone. Two-way RM ANOVA, time x inhibition, F_1,28_ = 7.447, *p = 0.0172. Sidak’s multiple comparisons test, mCh: Pre vs Post, ***p = 0.0003. hM4Di: Pre vs Post, p = 0.246. Right, direct comparison of CPA performance between mCh and hM4Di groups. Unpaired, two-tailed t-test, t_13_ = 2.81, *p = 0.0147. mCh, *n* = 7, hM4Di, *n* = 8. **o.** Left, mice with inhibition of *Pdyn* neurons fail to avoid the chamber paired with naloxone. Two-way RM ANOVA, time x inhibition, F_1,30_ = 0.0607, p = 0.809. Main effect of time, F_1,30_ = 33.09, ****p < 0.0001. Sidak’s multiple comparisons test, mCh: Pre vs Post, **p = 0.0016. hM4Di: Pre vs Post, **p = 0.0032. Right, direct comparison of CPA performance between mCh and hM4Di groups. Unpaired, two-tailed t-test, t_14_ = 0.985, p = 0.341. mCh, *n* = 8, hM4Di, *n* = 8.

To test the prediction that the NAc *Tshz1* neurons enriched with *Oprm1* are directly modulated by opioids, we infused GCaMP6f into the NAc medial shell of *Tshz1*-Flp mice or *Pdyn*-Cre mice, then administered escalating morphine doses prior to performing photometry recordings during acute naloxone withdrawal induction (Fig. 3e). *Tshz1* neurons exhibited decreased activity in response to morphine and a large rebound of increased activity above baseline in response to naloxone (Fig. 3f,g). The same mice treated with saline and naloxone showed no activity changes (Fig. 3f,h). In marked contrast, *Pdyn* neurons exhibited an inconsistent response to morphine, with a gradual decrease in average activity (Fig. 3i,j). Naloxone injection further decreased activity during withdrawal (Fig. 3i,j), and again no effects were observed with saline (Fig. 3i,k).

The differences in the activity patterns between NAc *Tshz1* or *Pdyn* neurons during withdrawal suggest that they play distinct behavioral roles. To test this prediction for withdrawal aversion, *Tshz1*-Flp and *Pdyn*-Cre mice were bilaterally injected with the inhibitory chemogenetic tool hM4Di into the NAc medial shell and conditioned in the withdrawal CPA assay after an injection of clozapine-*N*-oxide (CNO, 5 mg kg^-1^, ip) (Fig. 3l,m). Inhibition of *Tshz1* neurons blunted morphine withdrawal CPA learning (Fig. 3n), while inhibition of *Pdyn* neurons was ineffective (Fig. 3o). Based on our GCaMP6f recordings, we further predicted that activation of *Tshz1* neurons with the excitatory hM3Dq would interfere with morphine reward learning. While mCherry mice developed a strong morphine CPP, mice expressing hM3Dq did not (Extended Data Fig. 5a,b). Interestingly, excitation of *Tshz1* neurons with hM3Dq did not affect cocaine CPP (Extended Data Fig. 5c,d), suggesting specificity towards opioid reward. By contrast, inhibition of *Pdyn* neuron activity with Cre-dependent hM4Di had a marginal effect on morphine CPP (Extended Data Fig. 5e,f).

### *Tshz1* neurons modulate DA release through local inhibition in NAc

To explore the mechanisms by which NAc *Tshz1* neurons contribute to withdrawal aversion, we first determined the valence of NAc *Tshz1* neurons in a real time place preference assay in naïve mice. Stimulation of *Tshz1* neurons with channelrhodopsin (ChR2) led to an aversive experience as evidenced by avoidance of the stimulation-paired chamber (Fig. 4a,b). To determine if this aversion may be generated by decreases in DA release, which are commonly aversive^17–19^, we expressed GRAB DA in the NAc with Flp-dependent red-shifted ChRmine expressed only in *Tshz1* neurons (Fig. 4c). Stimulation of *Tshz1* neurons rapidly and robustly suppressed DA release in a highly repeatable fashion throughout each stimulation window (Fig. 4d-g). Furthermore, *Tshz1* neuron stimulation at different frequencies generated frequency-dependent increases in DA inhibition (Extended Data Fig. 6a-e). In contrast, stimulation of *Pdyn* neurons led to real time place preference (Extended Data Fig. 6f,g) and modest increases in DA release (Extended data fig. 6h-l).

**Fig 4.**
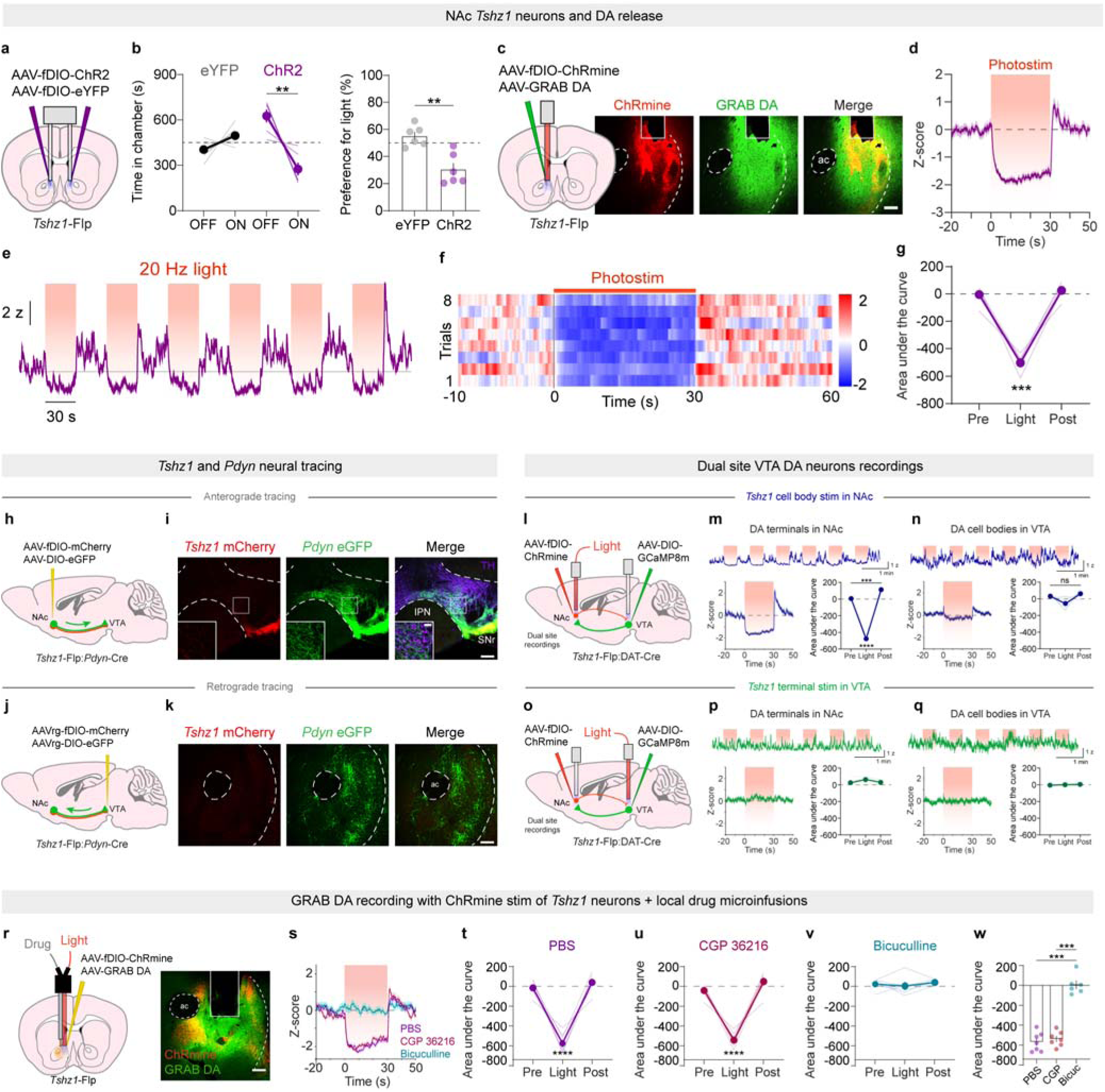
*Tshz1* neurons suppress DA release through local inhibition of DA terminals. **a.** Viral injections and fiber implants for real time place testing in *Tshz1*-Flp mice. **b.** Left, time spent in the light-paired chamber for each group. RM two-way ANOVA, time x light, F_1,10_ = 19.44, **p = 0.0013. Sidak’s multiple comparisons test, eYFP: OFF vs ON, p = 0.396. ChR2: OFF vs ON, **p = 0.0012. Right, direct comparison between groups. Unpaired, two-tailed t-test, t_10_ = 4.409, **p = 0.0013. eYFP, *n* = 6, ChR2, *n* = 6. **c.** Left, schematic of viral injection and fiber implant for simultaneous photometry recording and optical stimulation of NAc *Tshz1* neurons. Right, image of ChRmine and GRAB DA expression in the NAc. Scale = 100 um. **d.** Event histogram of average DA release while *Tshz1* neurons are stimulated with red light. **e.** Representative trace across multiple *Tshz1* neuron activation windows. **f.** Color map showing the effect of photomstimulation of *Tshz1* neurons on DA release. **g.** Area under the curve quantification of the peri-event histogram presented in panel **d**. RM one-way ANOVA, F_2,12_ = 92.34, ***p = 0.0004. Dunnett’s multiple comparison test, Pre vs Light, **p = 0.0017, Post vs Light, ***p = 0.0006, Pre vs Post, p = 0.2719. *n* = 5 mice. **h.** Anterograde tracing strategy in *Tshz1*-Flp:*Pdyn*-Cre mice. **i.** Image of NAc *Tshz1* and *Pdyn* inputs to the VTA. TH = tyrosine hydroxylase. Scale = 100 um. **j.** Retrograde tracing strategy in *Tshz1*-Flp:*Pdyn*-Cre mice. **k.** Image of retrograde labeled *Tshz1* and *Pdyn* neurons in the NAc. Scale = 100 um. **l.** Experimental strategy for simultaneous recordings of VTA DA cell bodies and axons while NAc *Tshz1* cell bodies are stimulated. **m.** Top, representative trace of DA axon activity across multiple stimulation windows. Bottom left, peri-event histogram of average DA terminal activity while *Tshz1* neurons are stimulated. Bottom right, area under the curve quantification. RM one-way ANOVA, F_2,18_ = 1739, ****p < 0.0001. Dunnett’s multiple comparison test, Pre vs Light, ****p < 0.0001, Post vs Light, ****p < 0.0001, Pre vs Post, ***p = 0.0002. *n* = 7 mice. **n.** Top, representative trace of DA cell body activity across multiple stimulation windows. Bottom left, peri-event histogram of average DA cell body activity while *Tshz1* neurons are stimulated. Bottom right, area under the curve quantification. RM one-way ANOVA, F_2,18_ = 7.460, *p = 0.0323. Dunnett’s multiple comparison test, Pre vs Light, p = 0.0648, Post vs Light, p = 0.0558, Pre vs Post, p = 0.0654. *n* = 7 mice. **o.** Experimental strategy for simultaneous recording of VTA DA cell bodies and axons while stimulating *Tshz1* terminals in the VTA. **p.** Top, representative trace of DA axons across multiple stimulation windows. Bottom left, peri-event histogram of average DA terminal activity while *Tshz1* terminals are stimulated. Bottom right, area under the curve quantification. RM one-way ANOVA, F_2,18_ = 3.358, p = 0.0809. Dunnett’s multiple comparison test, Pre vs Light, p = 0.156, Post vs Light, p = 0.1180, Pre vs Post, p = 0.9348. *n* = 7 mice. **q.** Top, representative trace of DA cell bodies across multiple stimulation windows. Bottom left, peri-event histogram of average DA cell body activity while *Tshz1* terminals are stimulated. Bottom right, area under the curve quantification. RM one-way ANOVA, F_2,18_ = 0.380, p = 0.6405. Dunnett’s multiple comparison test, Pre vs Light, p = 0.9132, Post vs Light, p = 0.8526, Pre vs Post, p = 0.4905. *n* = 7 mice. **r.** Configuration of GRAB DA photometry recording with drug microinfusions. Right, representative image of injection and recording site. Scale = 100 um. **s.** Peri-event histogram displaying average DA release during each drug condition. **t.** Area under the curve after PBS injection. RM one-way ANOVA, F_2,6_ = 81.96, ****p < 0.0001. Dunnett’s multiple comparison test, Pre vs Light, ***p = 0.0002, Post vs Light, ***p = 0.0001, Pre vs Post, p = 0.066. *n* = 7 mice. **u.** Area under the curve after CGP 36216 injection. RM one-way ANOVA, F_2,6_ = 89.44, ****p < 0.0001. Dunnett’s multiple comparison test, Pre vs Light, ****p < 0.0001, Post vs Light, ***p = 0.0001, Pre vs Post, *p = 0.0113. *n* = 7 mice. **v.** Area under the curve after Bicuculline injection. RM one-way ANOVA, F_2,5_ = 0.5157, p = 0.5409. Dunnett’s multiple comparison test, Pre vs Light, p = 0.8874, Post vs Light, p = 0.5501, Pre vs Post, p = 0.6213. *n* = 6 mice. **w.** Direct comparison of area under the curve for each drug condition. Ordinary one-way ANOVA, F_2,17_ = 81.83, ****p < 0.0001. Sidak’s multiple comparisons test, PBS vs CGP, p = 0.9033. PBS vs Bicuculline, ****p < 0.0001. CGP vs Bicuculline, ****p < 0.0001. *n* = 6-7 mice.

As canonical NAc D1 medium spiny neurons are known to project into the midbrain^38^, one hypothesis to explain the negative modulation of DA release by *Tshz1* neurons is that they directly inhibit DA neurons through projections to the VTA. To determine whether *Tshz1* or *Pdyn* neurons project to the VTA, we crossed *Tshz1*-Flp mice to *Pdyn*-Cre (*Tshz1*-Flp:*Pdyn*-Cre) mice and injected a cocktail of AAV-fDIO-mCherry and AAV-DIO-eGFP into the NAc (Fig. 4h). We observed robust *Pdyn* inputs to the VTA with only very sparse*Tshz1* inputs (Fig. 4i). Injection of retrograde AAVrg-fDIO-mCherry and AAVrg-DIO-eGFP into the VTA of separate *Tshz1*-Flp:*Pdyn*-Cre mice led to many eGFP retro-labeled *Pdyn* neurons in the NAc but very few mCherry *Tshz1* neurons; results that corroborate the anterograde tracing (Fig. 4j,k).

How might NAc *Tshz1* neurons exhibit a strong inhibitory influence over DA release with such minimal projections to the VTA? To test the hypothesis that these neurons modulate DA release locally in the NAc^39–43^, we crossed *Tshz1*-Flp mice to DAT-Cre (*Tshz1*-Flp:DAT-Cre) mice and expressed fDIO-ChRmine in the NAc and DIO-GCaMP8m in the VTA with optical fibers targeting both the NAc and ipsilateral VTA (Fig. 4l and Extended Data Fig. 7a,b). During simultaneous recordings, we stimulated *Tshz1* cell bodies via the fiber in the NAc (Fig. 4l) and observed a robust decrease in GCaMP8m activity in VTA DA axon terminals, but only minor changes in DA cell bodies (Fig. 4m,n). In the same mice, pulsing red light through the fiber in the VTA to activate *Tshz1* neuron VTA projections (Fig. 4o) had no effect on either VTA DA axon terminal or cell body activity (Fig. 4p,q). These results support the conclusion that *Tshz1* neurons modulate DA release in the NAc via local modulation of VTA DA axon terminals within the NAc.

To further examine this proposed functional anatomy, we injected *Tshz1*-Flp:DAT-Cre mice with AAV-fDIO-ChR2 into the NAc and AAV-DIO-tdTomato into the VTA to enable whole-cell recordings of identified DA neurons in response to optogenetic activation of *Tshz1* neuron axons (Extended Data Fig. 7d). We observed zero connectivity with DA neurons (Extended Data Fig. 8e-g; n=13 cells). To reduce sampling bias, we injected new *Tshz1*-Flp:DAT-Cre mice with a cocktail of AAV-fDIO-ChR2 and AAVrg-DIO-tdTomato into the NAc to selectively label NAc-projecting VTA DA neurons (Extended Data Fig. 7h). Whole-cell recordings from identified NAc-projecting DA neurons also yielded no detectable synaptic connectivity, with no cells (n=20 cells) responding to ChR2 stimulation of *Tshz1* inputs (Extended Data Fig. 7i-k).

To determine if *Tshz1* neurons modulate DA locally, we next injected *Tshz1*-Flp mice with fDIO-ChRmine and GRAB DA in the NAc and implanted a dual fiber-cannula implant for local drug microinfusions (Fig. 4r). Stimulation of *Tshz1* neurons produced the characteristic suppression of DA release after an injection of PBS (Fig. 4s,t). Microinfusion of the GABA_B_ receptor antagonist CGP 36216 also had no effect on the *Tshz1* neuron mediated depression of DA release (Fig. 4s,u). In contrast, the GABA_A_ receptor antagonist bicuculline completely abolished the effect of *Tshz1* neuron activation on DA release (Fig. 4s,v,w). These data support the hypothesis that the local inhibition of DA release in the NAc by *Tshz*1 neuron activation is due to local GABA release acting on GABA_A_ receptors.

### *Tshz1* promote withdrawal aversion through DA inhibition

Based on the strong expression of MORs in *Tshz1* neurons and their robust influence on DA release, we hypothesized that opioid withdrawal aversion requires activation of MORs in *Tshz1* neurons during opioid administration and rebound activation of these neurons during withdrawal leading to a decrease in DA. To test this hypothesis, we deleted *Oprm1* specifically in NAc *Tshz1* neurons by crossing *Tshz1*-Flp mice to floxed *Oprm1* (*Oprm1*^fl/fl^) mice and injecting Flp-dependent Cre-mCherry into the NAc (*Tshz1* MOR KO mice). A cohort of *Tshz1* MOR KO mice, in which we expressed GRAB DA in the NAc, were treated with morphine for 5 days, and then put into naloxone-precipitated withdrawal during DA photometry recordings (Fig. 5a). The initial increase in DA release resulting from morphine was unaffected compared to *Tshz1*-Flp:*Oprm1*^fl/fl^ mice that expressed mCherry in the NAc rather than Flp-dependent Cre-mCherry (Fig. 5b,c). However, MOR KO in *Tshz1* neurons significantly blunted the decrease in DA release evoked by naloxone (Fig. 5b,d). In addition, the frequency of transient DA release events was similar after morphine administration in the control and *Tshz1* MOR KO mice while the decrease in these events after naloxone was mostly prevented by deleting MORs from *Tshz1* neurons (Fig. 5e,f).

**Fig 5.**
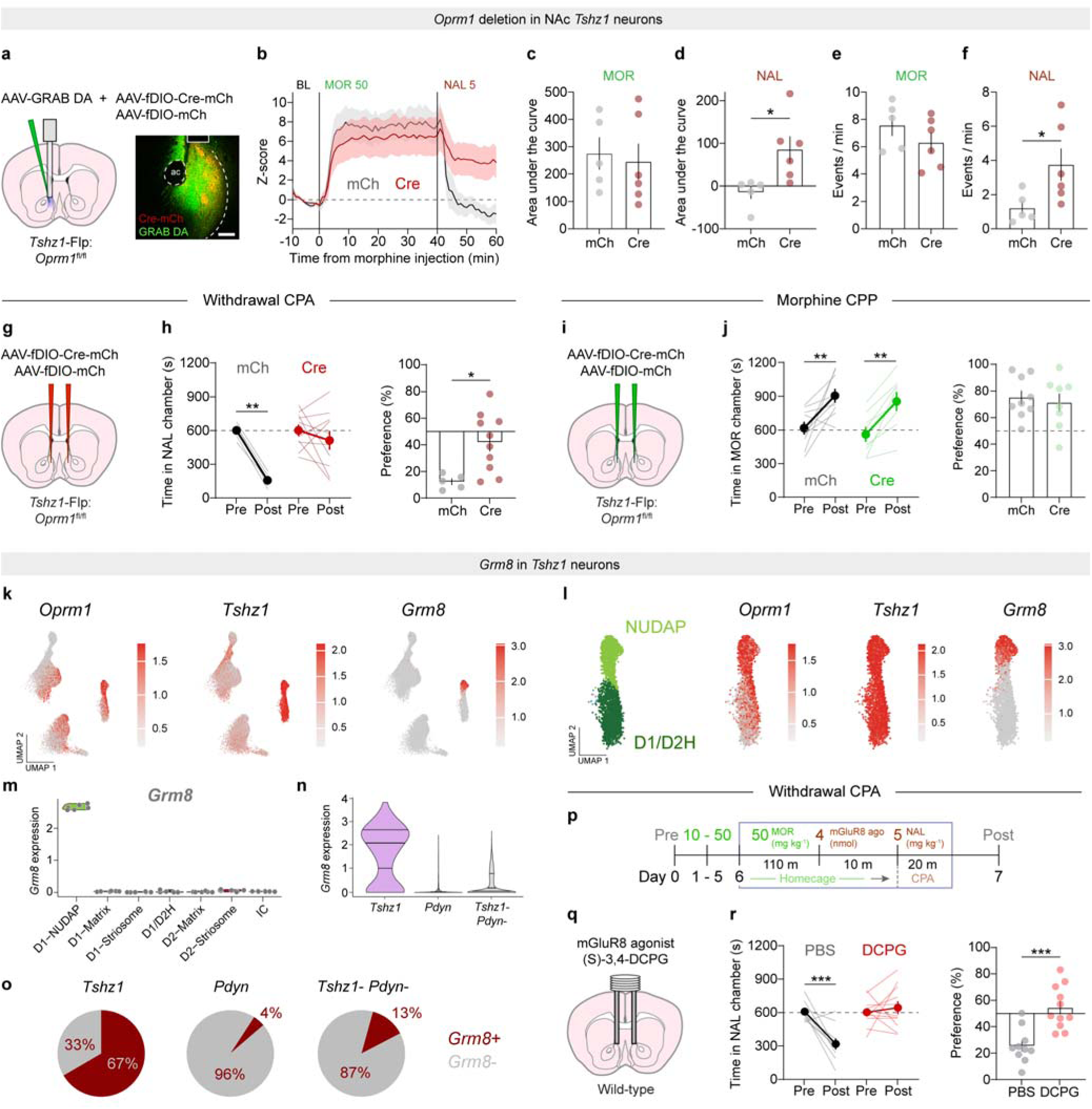
*Tshz1* neurons drive a hypodopaminergic state during withdrawal. **a.** Left, experimental strategy for selectively deleting *Oprm1* in NAc *Tshz1* neurons. Right, image of GRAB DA and Flp-dependent Cre-mCherry in NAc, scale = 100 um. **b.** Average bulk DA release in the NAc during morphine (50 mg kg^-1^) administration and precipitated withdrawal with naloxone (5 mg kg^-1^) in mice with deletion of *Oprm1* in NAc *Tshz1* neurons. **c.** Area under the curve during the morphine phase. Unpaired, two-tailed t-test, t_9_ = 0.328, p = 0.749. mCh, *n* = 5, Cre, *n* = 6. **d.** Area under the curve during the naloxone phase. Unpaired, two-tailed t-test, t_9_ = 2.735, *p = 0.023. mCh, *n* = 5, Cre, *n* = 6. **e.** DA release event frequency during the morphine phase. Unpaired, two-tailed t-test, t_9_ = 1.163, p = 0.2748. mCh, *n* = 5, Cre, *n* = 6. **f.** DA release event frequency during the naloxone phase. Unpaired, two-tailed t-test, t_9_ = 2.283, *p = 0.0484. mCh, *n* = 5, Cre, *n* = 6. **g.** Strategy for testing withdrawal CPA in mice with deletion of *Oprm1* in NAc *Tshz1* neurons. **h.** Left, time spent in the withdrawal-paired chamber during CPA. RM two-way ANOVA, time x deletion, F_1,13_ = 7.362, *p = 0.0177. Sidak’s multiple comparisons test, mCh: Pre vs Post, **p = 0.0022. Cre: Pre vs Post, p = 0.4503. Right, mice with *Oprm1* deletion avoid the withdrawal chamber significantly less than controls. Unpaired, two-tailed t-test, t_13_ = 2.91, *p = 0.0122. mCh, *n* = 6, Cre, *n* = 10. **i.** Strategy for testing morphine CPP in mice with deletion of *Oprm1* in NAc *Tshz1* neurons. **j.** Left, time spent in the morphine-paired chamber during CPP. RM two-way ANOVA, time x deletion, F_1,15_ = 0.0023, p = 0.9627. Main effect of time, F_1,15_ = 26.67, ***p = 0.0001. Sidak’s multiple comparisons test, mCh: Pre vs Post, **p = 0.0040. Cre: Pre vs Post, **p = 0.0054. Right, mice with *Oprm1* deletion avoid the withdrawal chamber significantly less than controls. Unpaired, two-tailed t-test, t_15_ = 0.4592, p = 0.6527. mCh, *n* = 9, Cre, *n* = 8. **k.** Feature plots showing the expression pattern of *Oprm1*, *Tshz1*, and *Grm8*. **l.** Expanded view of 3^rd^ cluster feature plots showing co-expression of *Oprm1*, *Tshz1*, and *Grm8* in the NUDAP archetype. **m.** Violin plot quantifying the expression pattern of *Grm8* in each cell archetype. **n.** Violin plot quantifying the expression pattern of *Grm8* in each cell-type. Mixed effect linear model, *Tshz1* (NUDAP) vs *Pdyn* ****p = 1.533e-15, *Tshz1* (NUDAP) vs *Tshz1*- *Pdyn*- ****p = 1.032e-90. *Tshz1* (NUDAP) vs *Pdyn,* Bonferroni corrected p = 3.066e-15. *Tshz1* (NUDAP) vs *Tshz1*- *Pdyn*-, Bonferroni corrected p = 2.065e-90. **o.** Pie charts illustrating the proportion of each cell-type expressing *Grm8*. **p.** Experimental timeline for CPA with mGluR8 agonist administration. **q.** Schematic of microinjection into NAc. **r.** Left, time spent in the withdrawal-paired chamber during CPA with mGluR8 activation. RM two-way ANOVA, time x drug, F1,19 = 12.97, **p = 0.0019. Sidak’s multiple comparisons test, PBS: Pre vs Post, ***p = 0.0007. DCPG: Pre vs Post, p = 0.7759. Right, mice with Grm8 agonist avoid the withdrawal chamber significantly less than controls. Unpaired, two-tailed t-test, t19 = 4.239, ***p = 0.0004. PBS, *n* = 10, DCPG, *n* = 11.

To test the behavioral relevance of these findings, we injected a new cohort of *Tshz1*-Flp:*Oprm1*^fl/fl^ mice with AAV-fDIO-Cre-mCherry or AAV-fDIO-mCherry bilaterally in the NAc and tested them for withdrawal CPA (Fig. 5g). Compared with mice expressing mCherry, *Tshz1* MOR KO mice displayed a significantly reduced aversion to the withdrawal-paired chamber (Fig. 5h). In a new cohort of mice, we then tested the emergence of physical withdrawal symptoms and did not detect any differences between control mice and those lacking MORs in *Tshz1* neurons (Extended Data Fig. 8a-d). In a final cohort, we tested morphine reward learning and observed nearly identical preference for the morphine-paired chamber in both lines of mice (Fig. 5i,j). These data indicate that MOR expression in NAc *Tshz1* neurons is required for the aversive component of withdrawal but not the associated physical symptoms nor for the rewarding property of morphine.

Since NAc *Tshz1* neurons are engaged during withdrawal and promote aversion, they offer a potential therapeutic target for reducing the aversion that is experienced during opioid withdrawal. Therefore, we searched our snRNAseq data for receptors preferentially expressed in *Tshz1* neurons and that would be expected to inhibit their activity. We identified *Grm8*, the type III inhibitory metabotropic glutamate receptor 8 (mGluR8), as a druggable target that is expressed almost exclusively in the D1-NUDAP archetype cluster, thereby co-localizing with *Oprm1* and *Tshz1* in mouse (Fig. 5k,l). We detected little *Grm8* in the other cell archetypes (Fig. 5m and Extended Data Fig. 8e-g), and also found that *Tshz1* NUDAP neurons were significantly more enriched with *Grm8* compared with *Pdyn* neurons or other remaining cell-types (Fig. 5n). Similarly, the proportion of *Tshz1* NUDAP neurons expressing *Grm8* was significantly larger than that for *Pdyn* or any other neurons (Fig. 5o). In human NAc, *GRM8* was also highly expressed in *TSHZ1* neurons, although its expression was less specific than observed in mouse (Extended Data Fig. 8h-o).

mGluR8 couples to a G_αi/o_ pathway and would be expected to inhibit neuronal activity^44,45^. Furthermore, agonists of mGluR8 are available and have been reported to reduce pain and anxiety^46,47^. Therefore we predicted that activating this receptor in the NAc would limit the increase in *Tshz1* neuron activity during withdrawal and thereby reduce the associated aversion. To test this hypothesis, we cannulated wild-type mice and microinjected the mGluR8 agonist (S)-3,4-DCPG into the NAc before withdrawal induction in the CPA assay (Fig. 5p,q). Strikingly, this treatment completely abolished withdrawal aversion learning compared to the robust CPA observed in mice that received intra-NAc infusion of PBS (Fig. 5r).

## Discussion

The opioid crisis has created a critical need for more effective interventions that break the vicious cycles of OUD^1-3,48^. Counteracting acute withdrawal distress could disrupt these cycles by preventing immediate, negatively reinforced relapse and slowing the transition to dependence. Effective interventions that target specific components of withdrawal require in depth knowledge of the underlying neural mechanisms, which have remained incompletely defined.

Here, we present a body of evidence that strongly supports the hypothesis that rapid suppression of DA release in the NAc is a critical component of acute withdrawal aversion and is largely mediated by a local NAc population of *Oprm1*-enriched cells defined by the expression of *Tshz1*. Surprisingly, *Tshz1* neurons regulate DA release locally in the NAc and not via influences on midbrain DA neuron activity. During administration of morphine, *Tshz1* neurons are inhibited but during precipitated withdrawal their activity increases resulting in a decrease in NAc DA release which mediates the aversion associated with withdrawal. Importantly, MOR-mediated influences on *Tshz1* neuron activity does not play any detectable role in the physical symptoms of withdrawal. Thus, we have defined a neural mechanism that clearly dissociates affective aversion from the physical signs of withdrawal. We also identify a molecular target, mGluR8, which is preferentially expressed in *Tshz1* neurons and the activation of which blocked withdrawal aversion. snRNAseq of human NAc tissue revealed the existence of analogous *Tshz1* neurons, suggesting that this neuronal population may be crucial for the development and maintenance of OUD in human subjects.

Neurons expressing *Tshz1* have been identified outside of the NAc although the functional significance of the transcription factor *Tshz1* expression is unclear. In the dorsal striatum, *Tshz1* neurons localize to striosomes where they respond to aversive stimuli and are essential to aversive learning^36^. Our snRNAseq analysis identified prototypical striosome genes in NAc *Tshz1* neurons (i.e., *Oprm1*, *Lypd1*, and *Nnat*)^34^ suggesting they may represent a ventral striatal striosome system. The intercalated cells of the amygdala also are enriched with *Tshz1*, *Oprm1*, and *Drd1* in mice, primates, and humans^49–51^. Functionally, they appear to gate amygdala outputs through feedforward inhibition to regulate fear and anxiety states^52–54^. This suggests that NAc *Tshz1* neurons may be part of a distributed network spanning the limbic forebrain and which plays critical roles in different forms of behavioral aversion.

Deletion of *Oprm1* specifically from *Tshz1* neurons prevented both opioid withdrawal aversion and reductions in DA release, but not physical withdrawal signs nor the reinforcement associated with morphine administration, which, consistent with a recent report^9^, we found required expression of MORs in midbrain GABA neurons. In contrast, the physical symptoms of withdrawal appear to require a distributed network of cells and circuits including the central amygdala^9^, thalamic inputs to the NAc^20^, striosomes^55^, prefrontal cortex^21^, and locus coeruleus^22–24^. Furthermore, the sociability deficits that occur during opioid withdrawal involve deficits in NAc 5-HT release due to kappa opioid receptor activity^56^. The mechanistic dissociation of the complex effects of opioids on the brain and behavior that contribute to OUD has important therapeutic implications. Indeed, our snRNAseq analysis identified *Grm8*, which has been identified in previous snRNAseq datasets^25,57^, as a preferentially expressed gene in *Oprm1*-enriched *Tshz1* neurons. Because mGluR8 functions as a type III inhibitory receptor, we tested whether its activation would ameliorate withdrawal aversion and found that direct injection of an mGluR8 agonist into the NAc attenuated withdrawal aversion learning, consistent with one previous study^58^.

Previous studies have reported that activation of cells in dorsal striatal striosomes can also reduce DA release^59,60^. A critical difference between NAc *Tshz1* neurons and those in striosomes is that the latter population projects strongly to the midbrain and influence DA release via actions on midbrain DA neurons whereas NAc *Tshz1* neurons send very sparse inputs to the VTA, which had no detectable effects on DA neuron cell body activity. Furthermore, our GCaMP8m recordings demonstrate strong modulation of local VTA DA axon activity in the NAc by *Tshz1* neurons. These effects contrast with NAc *Pdyn* neuron activation, which led to a potentiation of DA release and reinforcement^61^. Thus these two anatomically intermingled (adjacent) cell-types perform opposing actions on reward and reinforcement through their mechanistically distinct effects on DA release.

Alterations in the dynamics of striatal DA is a hallmark of drug dependent states^4,62^. Classic studies demonstrated clear increases in reward thresholds in subjects dependent on opioids^16,63,64^, suggesting suppression of the DA system. These observations are consistent with opponent process theories of addiction and implicate reduced DA function as a critical driver of drug seeking and relapse during withdrawal via negative reinforcement. Here, we show an unexpected mechanism of DA inhibition, NAc *Tshz1* neuron activation and local suppression of DA release during opioid withdrawal. Pharmacological experiments suggest a critical role for local GABA release acting on GABA_A_ receptors. Further work will be necessary to determine if this occurs via direct axo-axonic GABA synapses and/or through inhibition of NAc cholinergic interneurons, which can potentiate DA release^40,42^.

The ultimate challenge of treating OUD is preventing relapse the lifespan. We have identified NAc *Tshz1* neurons as a critical substrate for opioid withdrawal aversion, independent of physical symptoms. These findings add an important new component to the neural circuits that mediate the effects of opioids, while dissociating the distributed networks that control other important features of OUD. The discovery of these neurons also allowed for the identification of an unexpected molecular target, mGluR8, whose agonism may ameliorate aversion during withdrawal or opioid detoxification. The conservation of this cell-type in the human brain suggests that translation of these findings may have therapeutic implications for human OUD.

## Methods

### Subjects

Male and female C57BL/6J (Jackson Laboratory; stock #00664), *Pdyn*^tm1.1(cre)Mjkr^/LowlJ (*Pdyn*-IRES-Cre, Jackson Laboratory; stock #027958), *Tshz1*-2A-FlpO^36^ (a gift from Dr. Bo Li), *Slc6a3*^tm1.1(cre)bkmn^/J (DAT-Cre; Jackson Laboratory; stock #006660), *Oprm1*^tm1.1Cgrf^/KffJ (*Oprm1*^fl/fl^, Jackson Laboratory; stock #030074), and *Slc32a1*^tm2(cre)Lowl^/J (*Vgat*-IRES-Cre, Jackson Laboratory; stock #016962) mice were used. All mice (8-18 weeks old) were group housed on a 12-hr light/dark cycle with food and water *ad libitum*. All procedures compiled with animal care standards set forth by the National Institute of Health and were approved by the Stanford University’s Administrative Panel on Laboratory Animal Care and Administrative Panel of Biosafety.

### Viral vectors and stereotaxic surgery

AAVs purchased from Stanford Neuroscience Gene Vector and Virus Core: AAVdj-hSyn-Cre-eGFP, AAVdj-hSyn-eGFP, AAVdj-eF1a-fDIO-Cre-mCherry, AAVdj-eF1a-DIO-hChR2(H134R)-eYFP, AAVdj-eF1a-fDIO-hChR2(H134R)-eYFP, AAV8-Ef1a-DIO-rsChRmine-oScarlet, AAV8-Ef1a-fDIO-rsChRmine-oScarlet, AAVdj-eF1a-DIO-GCaMP6f, AAVdj-eF1a-fDIO-GCaMP6f, AAVdj-eF1a-DIO-hM4Di-mCherry, AAVdj-eF1a-fDIO-hM4Di-mCherry, AAVdj-eF1a-DIO-hM3Dq-mCherry, AAVdj-eF1a-fDIO-hM3Dq-mCherry, AAVdj-hSyn-DIO-eGFP, AAVdj-eF1a-DIO-mCherry, AAVdj-eF1a-fDIO-mCherry, AAVdj-eF1a-DIO-eYFP, AAVdj-eF1a-fDIO-eYFP, and AAVdj-Ef1a-DIO-tdTomato. AAVs purchased from Addgene: AAV9-hSyn-DIO-jGCaMP8m, AAVrg-hSyn-DIO-eGFP, AAVrg-hSyn-fDIO-mCherry, AAVrg-hSyn-DIO-tdTomato. AAV9-hSyn-GRAB-DA-2m (4.4) was purchased from WZ Biosciences (Columbia, MD). All viruses were injected 4-6 x 10^12^ infectious units per mL.

Mice of at least 7 weeks of age were anesthetized with isoflurane (1-2% v/v) and secured in a stereotaxic frame (David Kopf Instruments, Tujunga, CA). Viruses were injected into the NAc medial shell (AP +1.2; ML ±0.7; DV −3.6 from dura), NAc lateral core (AP +1.2; ML ±1.3; DV −3.4 from dura), VTA (AP −3.3; ML ±0.3; DV −4.2 from skull), or caudal VTA (AP −3.9; ML ±0.3; DV −4.2 from skull) at a rate of 150 nL min^-1^ (500 nL total volume) with a borosilicate pipette coupled to a pump-mounted 5 µL Hamilton syringe. Injector pipettes were slowly retracted after a 5 min diffusion period. For photometry and optogenetic experiments, mice were also implanted with optical fibers above the NAc (AP +1.2; ML ±0.7; DV −3.5 from dura), VTA (AP −3.3; ML ±0.4; DV −4.1 from skull), or caudal VTA (AP −3.9; ML ±0.3; DV −4.1 from skull). Optical fibers coupled to metal ferrules (Doric Lenses) with 400 µm core and 0.66 NA were unilaterally implanted over the NAc or midbrain for photometry recordings. Bilateral ‘guiding socket’ optical fibers (Doric Lenses) with 200 µm core and 0.66 NA were implanted in the NAc for optogenetic behavioral experiments. For photometry recordings with drug microinjections, a dual fluid injection cannula-optical fiber (OmFC; Doric Lenses) with 400 µm core and 0.66 NA was implanted over the NAc. For bilateral drug microinfusions, a bilateral guide cannula (P1 Technologies) was implanted over the NAc medial shell. Optical fibers and cannulas were secured to the skull with stainless steel screws (thread size 00-90 x 1/16, Antrin Miniature Specialties), C&B Metabond, and light-cured dental adhesive cement (Geristore A&B paste, DenMat). Mice were group housed to recover for at least 3 weeks before experiments began.

### Drug administration

To precipitate acute withdrawal, mice were made morphine dependent with 6 injections of 10-50 mg kg^-1^ morphine sulfate (Sigma Aldrich, M8777, ip) over 6 days in home cages^20^: i.e., 10 mg kg^-1^ on day 1, 20 mg kg^-1^ on day 2, etc. Mice received a second injection of 50 mg kg^-1^ morphine on day 6 before being injected with 5 mg kg^-1^ naloxone HCl (Tocris, 0599). During behavioral testing, mice were administered naloxone 2 hrs after the last morphine injection. During photometry recordings, mice were administered naloxone 40 min after the last morphine injection.

For local blockade of GABA receptors in the NAc during photometry, mice were prepared with the dual fluid injection cannula-optical fiber and habituated to the microinjection procedure 3 weeks later. CGP 36216 (3 mM in 300 nL, Tocris, 3219), 1(s),9(R)-(-)-Bicuculline methobromide (50 ng in 300 nL, Sigma Aldrich, 87561), or PBS was infused through a 100 um core polyimide injector cannula (FI_omFC-ZF_100/170) coupled to a 5 µL Hamilton syringe using a microinfusion pump (Harvard Apparatus) at a continuous rate of 100 nL min^-1^ to a total volume of 300 nL. Injector cannulas were removed 1 min after infusions were complete. Mice were allowed to recover for an additional 5 min before being connected to optical patch chords for rsChRmine stimulation recordings.

For stimulation of mGluR8 in the NAc, mice were prepared with bilateral guide cannulas targeting the NAc medial shell (P1 technologies). Mice were bilaterally injected with (s)-3,4-DCPG (4 nmol in 300 nL) at a continuous rate of 100 nL min^-1^ into the NAc. Injector cannulas were removed 1 min after infusions were complete. Mice were then allowed to recover for 10 min the homecage before being systemically injected with naloxone.

### Conditioned place preference

To evaluate positive reinforcing effects of morphine, mice were allowed to explore a 2-sided CPP chamber with distinct tactile floors and wall patterns (Med Associates Inc.) in a 20 min pretest. The next morning, mice were confined to one side of the chamber for 40 min after receiving an injection of saline. Approximately 4 hrs later, mice were confined to the opposite side of the chamber immediately after receiving an injection of morphine (20 mg kg^-1^, ip), again for 40 min. This procedure was repeated over the next 2 days for a total of 3 conditioning sessions. On the 5th day, mice were allowed to explore both sides of the CPP chamber for 20 min in a posttest. Preference was calculated as the percentage of time spent in the morphine-paired side of the chamber during the posttest. Morphine-paired sides were assigned in a counterbalanced and unbiased fashion such that the average preference for the morphine-paired side during the pretest was ∼50% for all groups.

Cocaine CPP was performed similarly to the procedure above with the following exceptions. Pretests, posttests, and conditioning sessions were 15 min. Mice received saline in the morning and 4 hrs later, mice were administered cocaine (15 mg kg^-1^, ip) and immediately confined to the appropriate side of the CPP chamber. Cocaine CPP was also performed in a counterbalanced and unbiased design.

For chemogenetic manipulations during CPP, mice were prepared with injections of hM4Di, hM3Dq, or mCherry and run through the CPP procedures about 4 weeks later. 30 min before each conditioning session, Clozapine-N-Oxide (CNO, 5 mg kg^-1^, ip) was injected in the home cage.

### Morphine withdrawal conditioned place aversion

To evaluate negative reinforcing effects of morphine, mice were allowed to explore a 2-sided CPP chamber with distinct tactile floors and wall patterns (Med Associates Inc.) in a 20 min pretest. Mice were then subjected to 5 injections of morphine (10-50 mg kg^-1^, ip) over 5 days in the home cage. On the 6^th^ day, mice were injected with one last dose of morphine (50 mg kg^-1^, ip) in the home cage and 120 min later, injected with naloxone (5 mg kg^-1^, ip) and confined to one side of the CPP chamber for a 20 min conditioning session. 24 hrs later, mice were allowed to explore both sides of the CPP chamber during a 20 min posttest. Preference was calculated as the percentage of time spent in the morphine-paired side of the chamber during the posttest. Naloxone-paired sides were assigned in a counterbalanced and unbiased fashion such that the average preference for the naloxone-paired side during the pretest was ∼50% for all groups.

For chemogenetic manipulations during CPA, mice were prepared with injections of hM4Di, hM3Dq, or mCherry and run through the CPA procedures about 4 weeks later. 30 min before each conditioning session, Clozapine-N-Oxide (CNO, 5 mg kg^-1^, ip) was injected in the home cage (i.e. 90 min after the morphine injection and 30 min before the naloxone injection).

For stimulation of mGluR8 in the NAc, mice were prepared with bilateral guide cannulas targeting NAc medial shell (P1 technologies). After habituation to microinjection procedures, mice were tested in the withdrawal CPA assay as described below. Before naloxone conditioning, mice were bilaterally injected with (s)-3,4-DCPG (4 nmol in 300 nL) into the NAc. Mice were then allowed to recover for 10 min in the homecage before systemic injection with naloxone and conditioned for CPA.

### Withdrawal physical signs

To determine the contribution of MORs in different populations to withdrawal physical signs, a group of *Oprm1*^fl/fl^ mice were bilaterally injected with AAV-Cre-eGFP or AAV-eGFP into the midbrain and a group of *Tshz1*-Flp:*Oprm1*^fl/fl^ mice were bilaterally injected with AAV-fDIO-Cre-mCherry or AAV-fDIO-mCherry into the NAc. Four weeks later, mice were subjected to the morphine dosing regimen outlined above in their home cages. On the sixth day, mice received a final injection of morphine (50 mg kg^-1^, ip) in the home cage. Two hrs later, naloxone (5 mg kg^-1^, ip) was administered and mice were immediately transferred to a 20 x 20 cm observation box with a camera mounted above for withdrawal recordings for 20 min. Withdrawal symptoms (jumping, rearing, and tremors) were carefully scored in 5 min bins, which were summed in the final analysis.

### Optogenetic stimulation

For behavioral optogenetic experiments, Prizmatix Pulsers and STSI LEDs were used to deliver blue light (450 nm, ∼10 mW, 5 ms pulses at 20 Hz) through a patch cord that mice were connected to via a rotary joint (Doric Lenses). For optogenetic stimulation during photometry recordings, Prizmatix Pulsers and STSI LEDs delivered red light (625 nm, 6 mW, 10 ms pulses at 1, 10, or 20 Hz) through a patch cord that connected to the same optical fiber used for the photometry recording via a rotary joint (Doric Lenses).

### Real-time place preference

The real-time place preference test was conducted in a rectangular box equipped with three chambers with similar floors and walls, separated by removable walls. Subjects were placed in the center chamber for 2 min at which point the barriers were lifted and the subject mouse was allowed to freely explore the entire apparatus for 15 min during which it received photostimulation with blue light (20 Hz, 10 ms pulses) whenever it entered the designated chamber, which was alternated between each testing session. After this initial phase, the side that was paired with photostimulation was swapped and mice were free to explore all three chambers for an additional 15 min reversal phase.

### Fiber photometry

Fiber photometry was performed as previously described^65^. Optical implants were connected to patch cables (400 um diameter, 0.57 numerical aperture (NA), Doric Lenses) via a ceramic sleeve (Doric).

Signals passed through a fiber optic commutator rotary joint (Doric) before filtering through a fluorescence mini cube (Doric) and reaching a femtowatt photodetector (2151, Newport). Frequency-modulated blue (465 nm) and UV (405 nm) LEDs (Doric) were used to stimulate GRAB DA and GCaMP emission and control signals through the same fibers. LED power was adjusted to ∼30 uW at the fiber tip. Digital signals were sampled at 1.0173 kHz, demodulated, lock-in amplified, and recorded by a real-time, lock-in signal processor (RZ5P, Tucker-Davis Technologies) using Synapse software (Tucker-Davis Technologies).

Signal processing was performed with custom scripts in MATLAB (MathWorks). Briefly, signals were down-sampled 10x and underwent Loess smoothing (window size = 30 ms). To minimize bleaching, we removed the first 5 min of each recording, when the steepest bleaching was likely to occur. For the full session analysis examining drug effects, 100 Hz down-sampled signals were de-bleached by fitting a mono-exponential decay function to the baseline (pre drug) portion of the signal and subtracting this fit curve from the full-length trace. To calculate dF/F, all fluorescence intensity values (F_465_) for the entire time course were referenced to the mean (F_mean_) fluorescence for all values of the de-bleached pre-drug baseline as (F_465_ - F_mean_) / F_mean_. Z-scoring was performed similarly, using only the pre-drug baseline to calculate F_mean_ and F_stdev_ (Z = [F_465_ – F_mean_] / F_stdev_). The resulting traces were smoothed (MATLAB, ‘filtfilt’) using a zero-phase moving average filter with an equally weighted 100 sample window. Transient detection was automated (MATLAB Signal Processing Toolbox, ‘findpeaks’), with a 3 sec lockout between events. Transients were defined as Z-scored fluorescence from −3 sec to +4 sec relative to the detected peak. Each transient was normalized to its baseline defined as −2.9 to −2 sec. For ChRmine stimulation recordings, the entire session was debleached with an iterative method which calculates dF/F in short moving windows, centers and normalizes these windows, and then repeats these calculations for 100 temporally-offset windows in the same session. The final debleached signal is the average of the 100 iterations.

ChRmine data were analyzed in 70 sec windows where red light stimulation occurred at time = 0 sec and lasted for 30 sec, resulting in 20 sec pre and post light periods. During drug administration recordings, the entire recording session was analyzed and signals were quantified for each drug phase. Z-scores of the fluorescence for each recording were calculated based on the F_mean_ and F_stdev_ of the local baseline signal before each event (−20 sec for ChRmine stimulation) or the baseline across the entire detrended session (−10 min for drug administration). To quantify bulk DA release or Ca^2+^ activity, area under the curve was calculated using trapezoidal numerical integration of the Z-scores for the windows defined for each experiment. Signals were aligned to the start of the optogenetic stimulation window or to each drug injection. For optogenetic stimulation, trials were combined into a single session average for each mouse and then averaged across mice. Peri-event histograms were constructed and area under the curve for each light or drug phase was calculated in MATLAB.

For DA recordings in the NAc, mice were injected with GRAB DA into the NAc medial shell with an optical fiber directed above. For simultaneous recordings of VTA DA cell bodies and terminals, DAT-Cre mice were injected with GCaMP8m into the VTA and implanted with two optical fibers targeting the VTA and ipsilateral NAc. To record caudal midbrain GABA neurons, *Vgat*-Cre mice were injected with GCaMP8m into the caudal VTA and implanted with an optical fiber over the same site.

To record NAc *Tshz1* and *Pdyn* neurons, *Tshz1*-Flp and *Pdyn*-Cre mice were injected with GCaMP6f in the NAc with fibers directed above. For photometry recordings during precipitated withdrawal, mice were prepared with viruses and optical fibers and allowed to recover for at least 3 weeks. Mice were then subjected to the morphine dependence regimen in the home cage for the first 5 days.

Recordings were performed on the 6^th^ day during an injection of morphine (50 mg kg^-1^, ip) followed by an injection of naloxone (5 mg kg^-1^, ip). On the test day, mice were connected to patch cables and allowed to habituate during a 10 min baseline recording period. Morphine was then injected and recordings continued for another 40 min. At this point, naloxone was injected and recordings continued for another 20 min.

For deletion of *Oprm1* specifically in *Tshz1* neurons, *Tshz1*-Flp mice were crossed to *Oprm1*^fl/fl^ mice and a 1:1 mixture of fDIO-Cre-mCherry (or fDIO-mCherry) and GRAB DA was injected into the NAc with a fiber above. These mice were brought through the same procedures as outlined above.

For ChRmine stimulation experiments, *Tshz1*-Flp or *Pdyn*-Cre mice were injected with a 1:1 cocktail of fDIO-ChRmine or DIO-ChRmine and GRAB DA in the NAc. A single optical fiber was implanted above the NAc for routing both blue light for photometry recordings and red light for ChRmine stimulation. ChRmine stimulation was achieved with Prizmatix Pulsers and STSI LEDs delivered red light (625 nm, 6 mW, 10 ms pulses at 1, 10, or 20 Hz) through a patch cord that connected to the same optical fiber used for the photometry recording via a rotary joint (Doric Lenses). ChRmine stimulation recordings consisted of a 10 min baseline recording period followed by ten 30-second windows every 30 seconds, for a 20 min recording. To determine the influence of NAc *Tshz1* neuron activation on the activity of VTA DA cell bodies and terminals, *Tshz1*-Flp mice were crossed to DAT-Cre mice and fDIO-ChRmine was injected into the NAc, DIO-GCaMP8m injected into the VTA, and optical fibers targeting the VTA and ipsilateral NAc were implanted. For examining pharmacology in the NAc during photometry, *Tshz1*-Flp mice were injected with the 1:1 cocktail of fDIO-ChRmine and GRAB DA and implanted with the dual fluid injection cannula-optical fiber. Mice that received drug microinjections did so 5 min before the 20 min recording.

To examine synchronized activity between VTA DA cell bodies and axons during withdrawal recordings, cross correlations were calculated with custom scripts in Python over a 5 sec time-lag window. The mean peak correlation was calculated for each drug phase of the recording, which included the 10 min baseline, the last 10 min of the morphine phase, and the first 10 min of the naloxone phase. Cross correlation plots and summary graphs were plotted in Prism.

### Single nuclei RNA sequencing and analysis

Single nuclei RNA seq data from nucleus accumbens of human and mouse were obtained from a previously reported multi-species atlas describing conserved medium spiny neuron (MSN) archetypes across primates and rodents^31^. Human data were sourced from multiple studies^31,57,66^ and mouse data from a single study^32^. Analysis was performed in R v4.4.1 using Seurat v5.0^67^ and associated packages.

Gene expression patterns were visualized using Uniform Manifold Approximation and Projection (UMAP) reduction with species-corrected effects, displaying both MSN archetype classifications and normalized gene expression (Fig. 2b,c,e,k,l; Fig. 5k,l; Extended Data Fig. 3b,g,I,n; Extended Data Fig. 8i,j). To reduce gene expression dropout while preserving cell type identity, metacell aggregation was applied using SuperCells R package v1.0^68^. This metacell reduction enabled quantification of relative gene expression relationships between *Tshz1*, *Pdyn*, and *Oprm1* in individual cells within each species (Fig. 2d,f,g,h,i,m,n; Fig. 5m,n,o; Extended Data Fig. 3c,d,e,f,j,k,l,m; Extended Data Fig. 8f,g,k,l,m,n,o). Expression thresholds for marker-positive neurons were empirically determined using the metacell-improved expression levels and used to calculate population distributions (Fig. 2i,n; Fig. 5o; Extended Data Fig. 8o).

Differential *Oprm1*/*OPRM1* expression between neuronal populations was assessed using linear mixed effects modeling (lme4 v1.1). For mouse data, the model *Oprm1 ∼ group + Sex + (1|Replicate)* was applied, where *group* term designated *Tshz1*+, *Pdyn*+, or double-negative populations, controlling for sex and repeated measures across biological replicates (Fig. 2h; Fig. 5n). For human data, the model *OPRM1 ∼ group + Sex + Project + (1|Replicate)* included an additional term to account for dataset-specific batch effects (Fig. 2m; Extended Data Fig. 8n).

Gene co-expression analysis was performed using voom normalization accounting for single cell heteroskedasticity and limma for differential expression testing^69,70^. Analysis included 19,317 genes exceeding minimum expression thresholds in mouse snRNA-seq metacells. A statistical model, *Y ∼ target_gene + sex + metacell_numcell*, was applied where *Y* represents all genes and *target_gene* represents *Tshz1*, *Pdyn*, or *Oprm1* expression within each metacell. Significantly co-expressed genes were identified using Bonferroni correction (p < 0.5/19,317) and filtered for positive correlations (log_2_(correlation) > 0; Fig. 2d,f,g). A similar analysis was used to identify pharmacological targets by identifying marker genes for NUDAP archetypes compared to any other neuron types. The list of marker genes was further filtered by cross referencing significant genes (Bonferroni adjusted p < 0.05) with the IUPHAR curated list of pharmacological targets^71^.

### Slice electrophysiology

For anterograde labeling of VTA neurons, adult female and male *Tshz1*-Flp:DAT-Cre mice were injected with AAVdj-Ef1a-DIO-tdTomato into the VTA and AAV-fDIO-ChR2-eYFP into the NAc. For retrograde labeling of VTA neurons, adult female and male *Tshz1*-Flp:DAT-Cre mice were injected with a 1:1 cocktail of AAVrg-hSyn-DIO-tdTomato and AAV-fDIO-ChR2-eYFP into the NAc. Acute coronal brain slices containing the VTA were made 4-6 weeks after viral injection for recordings.

Briefly, mice were anesthetized with isoflurane and transcardially-perfused with ice-cold oxygenated artificial cerebrospinal fluid (ACSF, in mM): 126 NaCl, 21.4 NaHCO_3_, 2.5 KCl, 1.2 NaH_2_PO_4_, 2.4 CaCl_2_, 1.0 MgSO_4_, 11.1 glucose and 5 sodium ascorbate. Following perfusion, brains were rapidly dissected and coronal 220 μM sections were cut at 0.1mm/min on Leica 1200-S in oxygenated ACSF + sodium ascorbate and transferred to oxygenated HEPES solution (in mM): 86 NaCl, 2.5 KCl, 1.2 NaH_2_PO_4_, 35 NaHCO_3_, 20 HEPES, 25 glucose, 5 sodium ascorbate, 2 thiourea, 3 sodium pyruvate, 1 MgSO_4_-7H_2_O, 2 CaCl_2_-2H_2_O for 10 minutes in 34°C water bath. Slices were then sustained at room temperature in HEPES solution until recording.

Slices were then transferred to a recording chamber where they were submerged in ACSF without sodium ascorbate, at 30°C with a flow rate of 1-2 mL/min. ACSF included 100 μM DNQX to block AMPAR-mediated currents throughout the recordings. VTA neurons were visualized using DIC and neurons were selected for patching by visually identifying the presence of tdTomato fluorescence. We sampled VTA dopamine neurons along the anterior-posterior axis with bias towards the ventro-medial VTA where the maximum *Tshz1* innervation was observed. Patch pipettes were filled with (in mM): 125 KCl, 2.8 NaCl, 5 MgCl_2_, 0.2 CaCl_2_, 2 Na^+^-ATP, 0.3 Na^+^-GTP, 0.6 EGTA, and 10 HEPES.

Voltage clamp recordings were performed at −70 mV and series resistance was continuously monitored with a 5 mV hyperpolarizing step. Clampex software triggered two 5 ms pulses of blue light (472 nm LED, Thorlabs) separated by 50 ms through the 40x water immersion objective. Sweeps were performed every 30 sec and a minimum of 8 sweeps were collected to identify a light-evoked input. Recordings with changes in input resistance or series resistance greater than 10% were discarded and not used in analysis. Holding current was monitored throughout and recordings with significant deviations in holding current or baseline holding currents greater than 500 pA were not used. All electrophysiology data was analyzed using Clampfit software.

### Fluorescence *in situ* hybridization

Wild-type brains were flash frozen in isopentane on dry ice, sectioned on a cryostat at 16 µm, and processed for fluorescence *in situ* hybridization (FISH) with the RNAscope Multiplex Fluorescent v2 assay according to the manufacturer’s guidelines (Advanced Cell Diagnostics). Transcripts examined were *Tshz1* (ACDBio cat# 494291), *Pdyn* (ACDBio cat# 318771), and *Oprm1* (ACDBio cat# 315841). Slides were coverslipped with Fluoromount-G with DAPI (Southern Biotech, 0100-20) and stored at 4°C in the dark before imaging. To prevent over-digestion, the RNAscope protocol was modified with 1 hr of 4% PFA fixation and the application of protease plus for 15 min.

Stitched images were collected on a Keyence BZ-X800 microscope with a 10x, 0.45 NA objective at a resolution of 0.755 um/pixel, with identical exposure settings for all samples, and image analysis was done in FIJI. Images were first background subtracted across all channels (50 pixel rolling ball radius) and then the channels were split for further processing. The *Oprm1*, *Tshz1*, and *Pdyn* channels were autothresholded using the Triangle procedure. The DAPI channel of each image was used for cell detection and segmentation. This was accomplished by preprocessing the images with a top hat filter (radius 12 pixels), a sharpening operation, and then a second top hat filter (7 pixels). After this, the images were auto-thresholded with Li’s algorithm, followed by an opening operation and watershedding. Nuclei were then detected using the Analyze Particles function (size=64-324 pixels, circularity=0.25-1.00). The ROI manager was then used to measure the brightness and percentage of area covered by signal in each of the *Oprm1*, *Tshz1*, and *Pdyn* channels for each detected cell. For statistical comparisons, data were averaged across cells within animals.

### Histology

Mice were anesthetized with isoflurane and perfused transcardially with 1X PBS followed by 4% paraformaldehyde in PBS, pH 7.4. Brains were extracted and post-fixed overnight in the same fixative. Brains were sectioned at 40 µm on a vibratome and collected in PBS. Free-floating sections were washed three times in PBS with 0.2% Triton X-100 (PBST) for 10 min at room temperature and then incubated in a blocking solution containing PBST with 3% normal goat serum for 1 hr. Sections were next incubated in mouse anti-TH (1:1000, Immunostar, 22941), rabbit anti-RFP (1:1000, Rockland, 600-401-379), or chicken anti-GFP (1:1000, Aves Labs, GFP-1010) in blocking solution rotating at 4°C for 24 hr. After three 10 min washes in PBST, sections were incubated in species-specific secondary antibodies Alexa Fluor 488, 594, or 647 (1:700, Invitrogen, A-11039, A-11058, and A-31573) in blocking solution for 1 hr at room temperature. Finally, sections were washed three times for 10 min in PBS, mounted onto SuperFrost Plus glass slides, and coverslipped with Fluoromount-G with DAPI (Southern Biotech, Birmingham, AL, 0100-20). Fluorescent images were collected on a Nikon A1 confocal microscope or a Keyence BZ-X800 fluorescence microscope.

### Statistical analysis

Investigators were blinded to the manipulations that experimental subjects had received during behavioral testing, recordings, and data analysis. All behavioral data were analyzed and graphed with GraphPad Prism 9. All photometry data were processed and analyzed in MATLAB. Cross correlation analysis was performed in Python. All transcriptomic data were processed and analyzed in RStudio. Data distribution and variance were tested using Shapiro-Wilk normality tests. Normally distributed data were analyzed by unpaired, two-tailed t-tests, or one- or two-factor repeated measures ANOVA with *post-hoc* Sidak’s, Tukey’s, or Dunnett’s correction for multiple comparisons. When normal distributions were not assumed, the Wilcoxon signed rank test was performed for within group comparisons of two treatments, Mann-Whitney for between group comparisons, and Kruskal-Wallis with post-hoc Dunn’s test for multiple comparisons. Paired comparisons were performed when appropriate. Differences were considered significant when p < 0.05. All pooled data are expressed as mean ± SEM.

*References 65-71 are Methods-only.

## Acknowledgements

We thank the Malenka Lab, Eshel Lab, Heifets Lab, Rajani Maiya, and Paul Kramer for critical discussion. We thank the Stanford Gene Vector and Virus core for reagents. We thank Dr. Bo Li for sharing *Tshz1*-2A-FlpO mice. This work was supported by philanthropic funds donated to the Nancy Pritzker Laboratory at Stanford University. M.B.P. was supported by NIH grant K99 DA056573. J.M.T. was supported by NIH grant K08 DA055157, a Brain and Behavior Research Foundation Young Investigator Grant, and the Stanford University School of Medicine Department of Psychiatry & Behavioral Sciences 2024 Innovator Grants Program. D.F.C.P. was supported by an NSF Graduate Research Fellowship and an HHMI Gilliam Fellowship for Advanced Study. B.N.P. was supported by NIH grant F30 DA053020. Z.F. was supported by NIH grants R01 DA061243, R01 ES034037, R01 DK124219, R21 DA052419, the Baszucki Group, and The Pittsburgh Foundation. A.R.P. was funded by NIH grants UF1 MH130881 and DP1 DA046585. N.E. was supported by NIH grant K08 MH123791, a Brain & Behavior Research Foundation Young Investigator Grant, a Burroughs Wellcome Fund Career Award for Medical Scientists, and a Simons Foundation Bridge to Independence Award. R.C.M. was supported by a grant from the Stanford Wu Tsai Neurosciences Institute, a grant from the UCSF Dolby Family Center for Mood Disorders, and NIH grant P50 DA042012.

## Author contributions

M.B.P., J.M.T., and R.C.M. conceived the study and designed the experiments. Fiber photometry experiments were performed by M.B.P., G.C.T., J.M.T., and A.S. Optogenetic experiments were performed by M.B.P. and G.C.T. Behavioral experiments were performed by M.B.P., J.M.T. and J.A.G.S. Electrophysiology recordings were performed by N.D. and R.S. FISH experiments were performed by D.F.C.P. and M.G. Programming and data analysis were performed by J.B and Z.Z. snRNAseq analysis was performed by B.N.P. with supervision from A.P. and Z.F. Anatomical tracing was performed by M.B.P and J.M.T. Histological analysis was performed by J.A.G.S. and J.M.T. A.P.F.C. assisted with surgical procedures. N.E. provided critical hardware, wrote critical analysis software, and contributed to data interpretation. The manuscript was written by M.B.P., J.M.T., and R.C.M. and edited by all authors.

## Data availability

The datasets generated and analyzed in this study are available from the corresponding authors upon reasonable request.

## Code availability

No new single cell or nuclei RNA sequencing experiments were generated for this study. All human and mouse single cell data and annotations were accessed from the multi-species striatum atlas as reported by Gayden et al^31^. The analysis scripts used to create the main and supplemental figures for this study will be deposited online at the GitHub repository https://github.com/pfenninglab/StriatumComparativeGenomics/. Any intermediate files not available here can be requested from the corresponding author. Code used for data processing and analysis is available from the corresponding authors upon reasonable request.

## Extended Data Figure

**Extended Data Fig 1.**
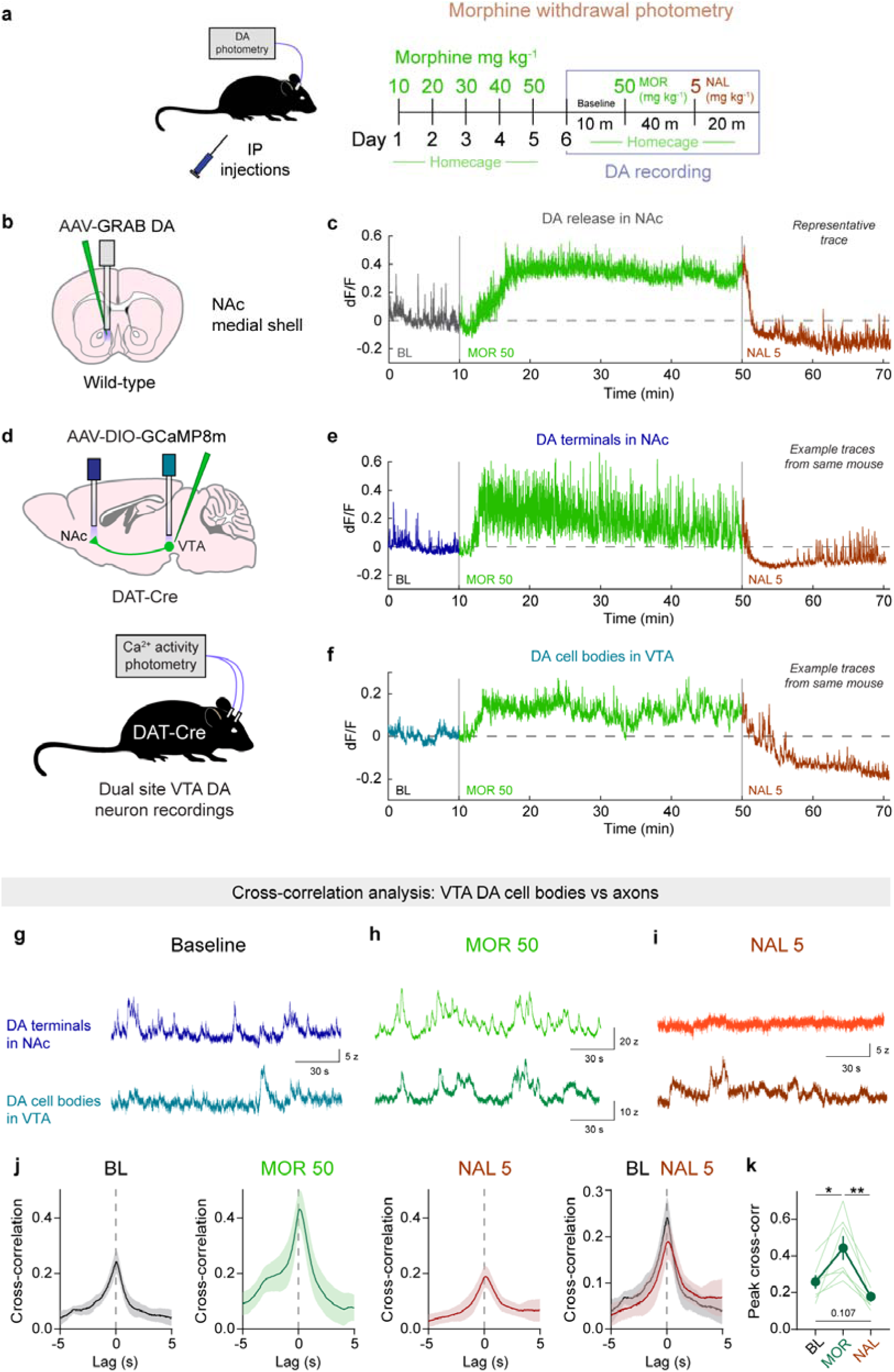
Further analysis of withdrawal-induced suppression of DA release in the NAc. **a.** Schematic of DA photometry recording and experimental schedule. **b.** Schematic of DA recoding in NAc medial shell of wild-type mice. **c.** Full session withdrawal recording trace of DA release from a representative mouse. **d.** Top, schematic of dual calcium recordings in NAc and VTA. Bottom, schematic of DAT-Cre mouse with the dual site VTA DA neuron recording. **e.** Full session withdrawal recording trace of DA terminals in NAc from a representative mouse. **f.** Full session withdrawal recording trace of DA cell bodies in VTA from the same representative mouse presented in panel **e**. **g.** Example traces from simultaneous recordings of VTA DA terminals in NAc and cell bodies in VTA during the baseline period in a representative mouse. **h.** Example traces from simultaneous recordings of VTA DA terminals in NAc and cell bodies in VTA during the morphine (MOR 50) phase in a representative mouse. **i.** Example traces from simultaneous recordings of VTA DA terminals in NAc and cell bodies in VTA during the naloxone (NAL 5) phase in a representative mouse. **j.** Average cross-correlation between simultaneous traces from VTA DA terminals and cell bodies for each recording phase. Left, baseline (BL). Center left, morphine (MOR 50). Center right, naloxone (NAL 5). Right, direct comparison of BL and NAL 5 phases. **k.** Summary graph of average peak cross-correlation for each drug phase. RM one-way ANOVA, F_2,19_ = 14.43, **p = 0.0024. Tukey’s multiple comparisons test, BL vs MOR *p = 0.0482, MOR vs NAL, **p = 0.0072, BL vs NAL p = 0.107.

**Extended Data Fig 2.**
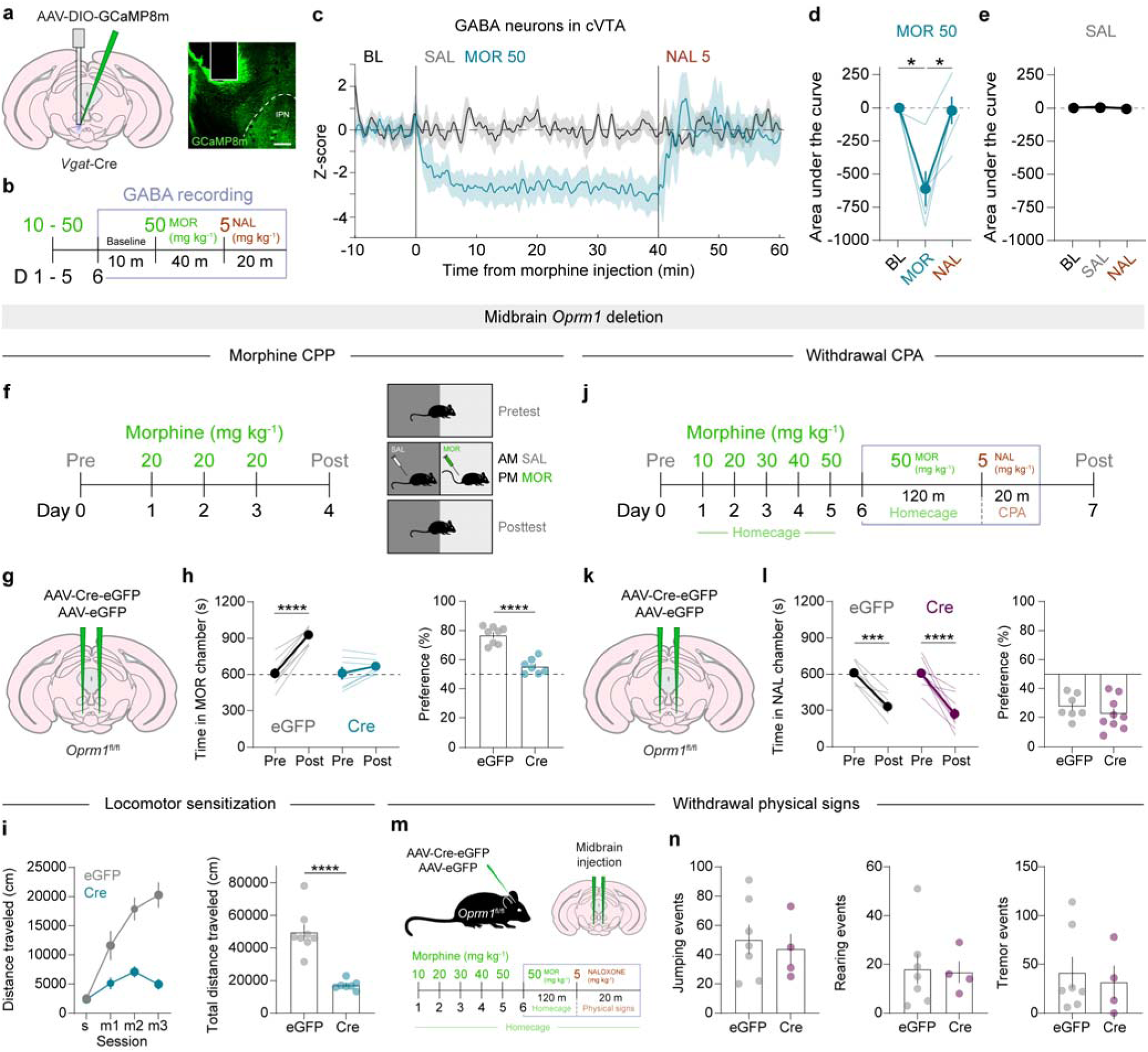
Midbrain GABA neurons mediate opioid reward, but not withdrawal. **a.** Schematic of calcium recordings in caudal midbrain GABA neurons during precipitated opioid withdrawal. **b.** Time course of experimental design. **c.** Bulk activity of midbrain GABA neurons during morphine (50 mg kg^-1^) administration and precipitated withdrawal with naloxone (5 mg kg^-1^). *n* = 5 mice. **d.** Area under the curve for each drug phase. RM one-way ANOVA, F_2,12_ = 15.44, **p = 0.0027. Dunnett’s multiple comparisons test, BL vs MOR *p = 0.0204, MOR vs NAL, *p = 0.0168, BL vs NAL p = 0.9704. **e.** Area under the curve for each drug phase after saline injection. RM one-way ANOVA, F_2,7_ = 0.451, p = 0.5792. *n* = 3 mice. **f.** Experimental design and time course of morphine conditioned place preference (CPP) test. **g.** Schematic of bilateral deletion of *Oprm1* in the caudal midbrain. **h.** Left, time spent in morphine-paired chamber pre and post conditioning. RM two-way ANOVA, time x deletion F_1,13_ = 34.64, ****p < 0.0001. Sidak’s multiple comparisons test, eGFP: Pre vs Post, ****p < 0.0001. Cre: Pre vs Post, p = 0.191. Right, direct comparison of CPP performance. Two-tailed, unpaired t-test, t_13_ = 7.458, ****p < 0.0001. eGFP, *n* = 8, Cre, *n* = 7. **i.** Left, time course of behavioral sensitization to morphine. RM two-way ANOVA, time x drug, F_1,58_ = 9.579, ****p < 0.0001. Sidak’s multiple comparisons test, eGFP vs Cre; SAL day 1 p = 0.9986, MOR day 1 p = 0.1490, MOR day 2 **p = 0.0023, MOR day 3 ***p = 0.0003. Right, comparison of total distance traveled over 3 days over morphine conditioning. Two-tailed, unpaired t-test, t_13_ = 6.248, ****p < 0.0001. eGFP, *n* = 8, Cre, *n* = 7. **j.** Experimental design and time course of morphine withdrawal conditioned place aversion (CPA) test. **k.** Schematic of bilateral deletion of *Oprm1* in the caudal midbrain. **l.** Left, time spent in withdrawal-paired chamber pre and post conditioning. RM two-way ANOVA, time x deletion, F_1,14_ = 0.564, p = 0.465. Main effect of time, F_1,14_ = 63.62, ****p < 0.0001. Sidak’s multiple comparisons test, eGFP: Pre vs Post, ***p < 0.0005. Cre: Pre vs Post, ****p < 0.0001. Right, direct comparison of CPP performance. Two-tailed, unpaired t-test, t_14_ = 0.997, p = 0.336. eGFP, *n* = 7, Cre, *n* = 9. **m.** Experimental design of withdrawal physical sign testing. **n.** Quantification of physical signs during precipitated withdrawal. Jumping events, two-tailed, unpaired t-test, t_9_ = 0.4305, p = 0.6770. Rearing events, two-tailed, unpaired t-test, t_9_ = 0.1559, p = 0.8795. Tremor events, two-tailed, unpaired t-test, t_9_ = 0.3904, p = 0.7054. eGFP, *n* = 7, Cre, *n* = 4.

**Extended Data Fig 3.**
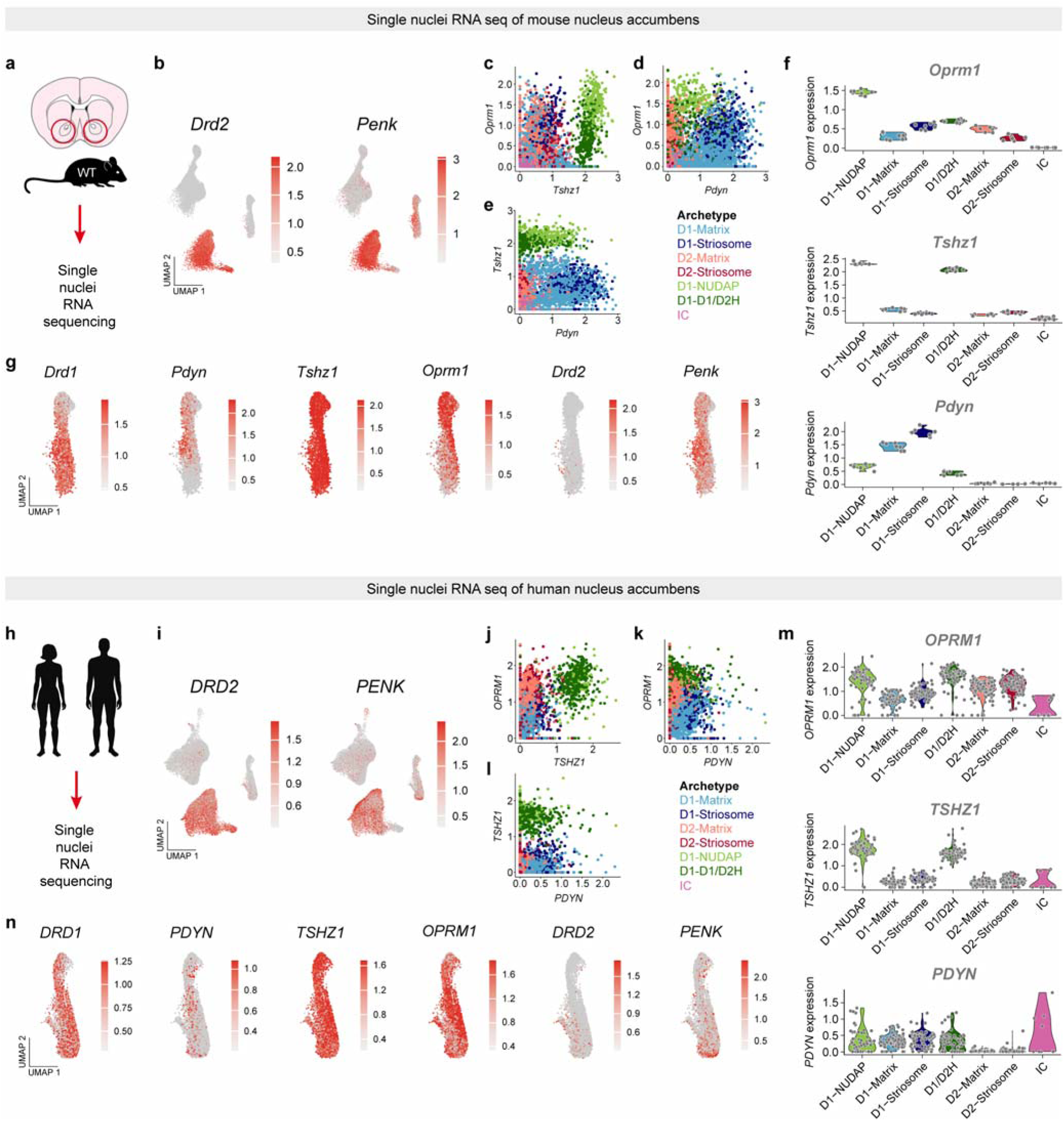
Additional snRNAseq data. **a.** Single nuclei RNAseq of mouse NAc. **b.** Feature plots showing the expression patterns of *Drd2* and *Penk*. **c.** Plot showing relationship between *Tshz1* and *Oprm1* expression in all cells colored by archetype. D1-NUDAP cells stand out as containing high expression of both *Tshz1* and *Oprm1*. **d.** Plot showing relationship between *Pdyn* and *Oprm1* expression in all cells colored by archetype. **e.** Plot showing relationship between *Pdyn* and *Tshz1* expression in all cells colored by archetype. **f.** Violin plots showing the expression of *Oprm1* (top), *Tshz1* (middle), and *Pdyn* (bottom) in each archetype. **g.** Feature plots showing key genes in an expanded view of the eccentric cluster. **h.** Experimental design of single nuclei RNAseq of human NAc. **i.** Feature plots showing the expression patterns of *DRD2* and *PENK*. **j.** Plot showing relationship between *TSHZ1* and *OPRM1* expression in all cells colored by archetype. D1-NUDAP and D1-D1/D2 cells stand out as containing high expression of both *TSHZ1* and *OPRM1*. **k.** Plot showing relationship between *PDYN* and *OPRM1* expression in all cells colored by archetype. **l.** Plot showing relationship between *PDYN* and *TSHZ1* expression in all cells colored by archetype. **m.** Violin plots showing the expression of *OPRM1* (top), *TSHZ1* (middle), and *PDYN* (bottom) in each archetype. **n.** Feature plots showing key genes in an expanded view of the eccentric cluster.

**Extended Data Fig 4.**
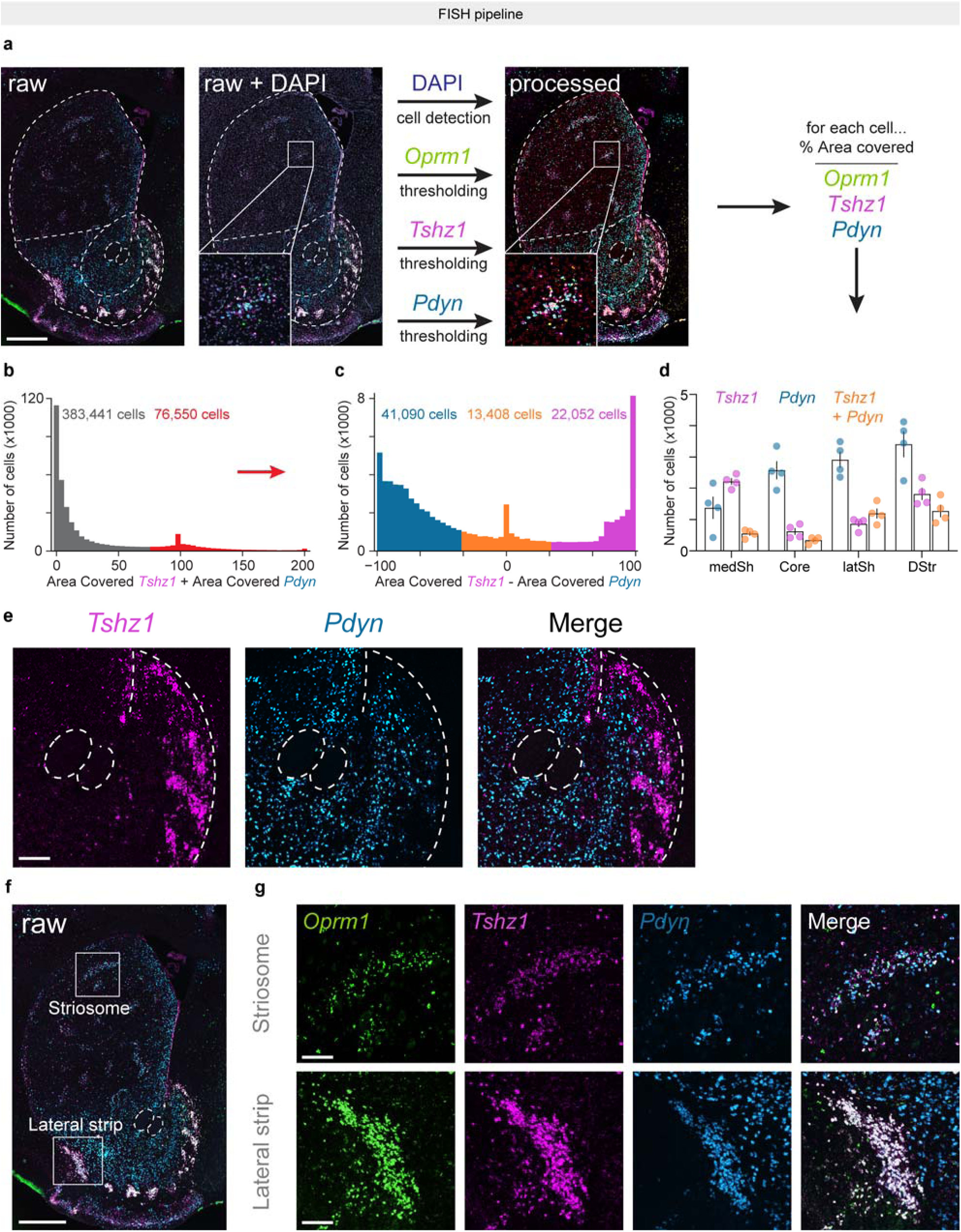
FISH pipeline and additional anatomical data. **a.** Illustration of FISH pipeline with RNAscope. Raw image data is merged with DAPI. Image processing undergoes cell detection with DAPI mask, then all 3 fluorescence channels are thresholded and the remaining signal for each channel is quantified within each detected cell. Inset shows an example striosome in the dorsal striatum. Red outlines in the processed images display detected cells. Scale = 500 um. **b.** All detected cells (459,991 cells) were analyzed for expression of *Tshz1* and *Pdyn*. Threshold for cells expressing at least one gene was set to 75 area units covered. A minority of detected cells across the entire striatum (76,550 / 459,991 = 16.6%) was positive for *Tshz1* or *Pdyn* and used in the analysis. **c.** Positive cells identified in panel b were further analyzed for co-localization of *Tshz1* and *Pdyn*, and classified as either *Tshz1*+*, Pdyn*+, or *Tshz1* + *Pdyn* based on a simple subtraction algorithm (*Tshz1* area covered – *Pdyn* area covered). Cells with a threshold of +/-25 area units covered by both genes were classified as co-expressing cells. **d.** Quantification of all three cell-types identified in each striatal subregion. *n* = 4 mice. **e.** Image demonstrating minor overlap of *Tshz1* and *Pdyn* in the NAc medial shell. Scale = 100 um. **f.** Raw image with regions of interest featuring striosomes in the dorsal striatum and the lateral strip in the lateral ventral striatum. Scale = 500 um. **g.** Top, image of striosomes featured in the region of interest in panel f. Presence of all three genes is clear. Bottom, image of the lateral strip featured in the region of interest in panel f. Strong co-localization of all three genes is clear. Scale = 100 um.

**Extended Data Fig 5.**
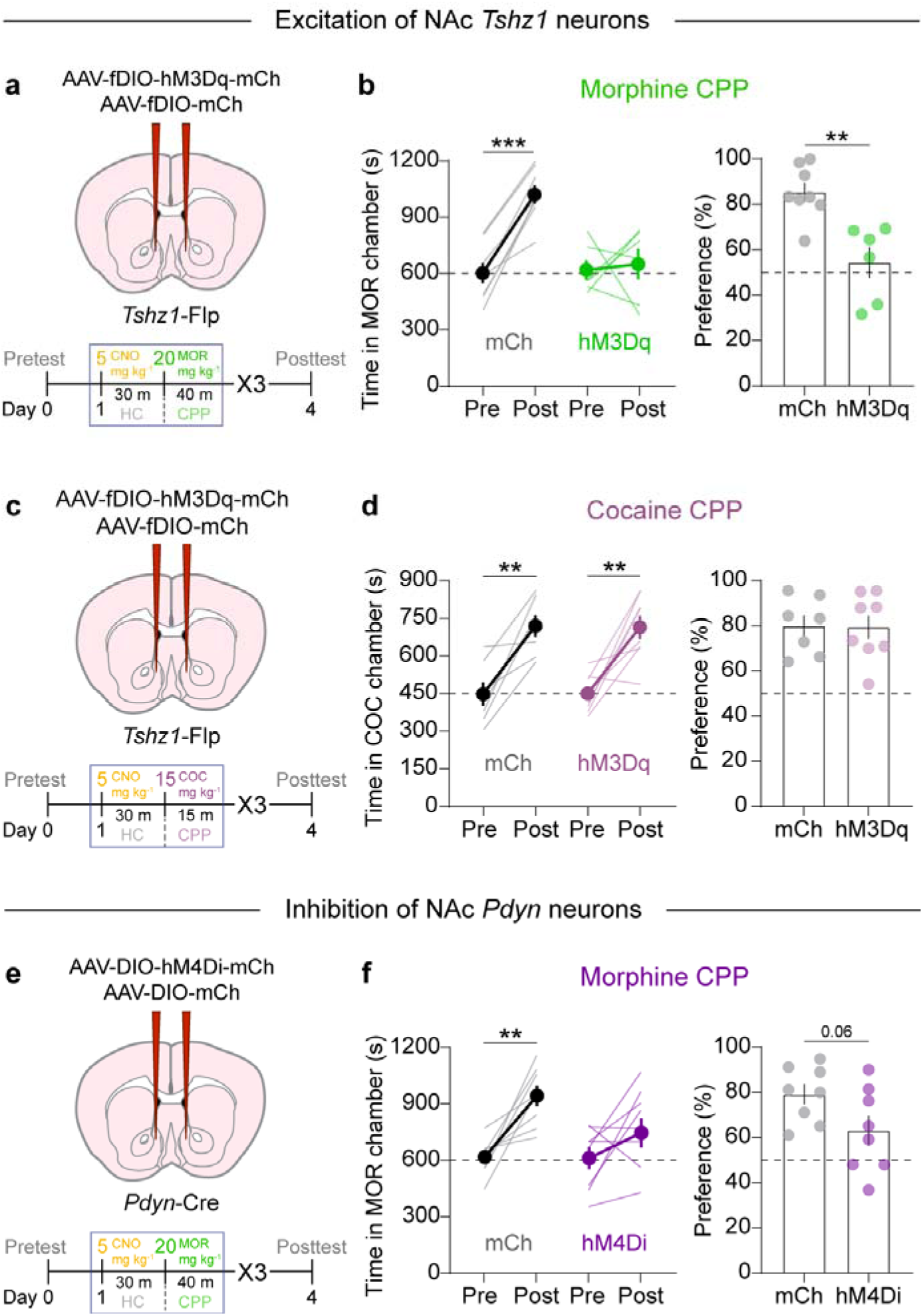
Additional CPP data from chemogenetic manipulation of NAc *Tshz1* and *Pdyn* neurons. **a.** Schematic of experimental design and time course of morphine CPP with excitation of *Tshz1* neurons. **b.** Left, time spent in morphine-paired chamber pre and post conditioning. RM two-way ANOVA, time x excitation F_1,26_ = 11.36, **p = 0.0056. Sidak’s multiple comparisons test, mCh: Pre vs Post, ***p = 0.0002. hM3Dq: Pre vs Post, p = 0.924. Right, direct comparison of CPP performance. Two-tailed, unpaired t-test, t_12_ = 4.116, **p = 0.0014. mCh, *n* = 8, hM3Dq, *n* = 6. **c.** Schematic of experimental design and time course of cocaine CPP with excitation of *Tshz1* neurons. **d.** Left, time spent in cocaine-paired chamber pre and post conditioning. RM two-way ANOVA, time x excitation F_1,28_ = 0.0075, p = 0.9322. Main effect of time F_1,28_ = 38.76, ****p < 0.0001. Sidak’s multiple comparisons test, mCh: Pre vs Post, **p = 0.0017. hM3Dq: Pre vs Post, **p = 0.0012. Right, direct comparison of CPP performance. Two-tailed, unpaired t-test, t_13_ = 0.085, p = 0.933. mCh, *n* = 7, hM3Dq, *n* = 8. **e.** Schematic of experimental design and time course of morphine CPP with inhibition of *Pdyn* neurons. **f.** Left, time spent in morphine-paired chamber pre and post conditioning. RM two-way ANOVA, time x excitation F_1,30_ = 0.1736, p = 0.2089. Main effect of time F_1,30_ = 18.1, ***p = 0.0008. Sidak’s multiple comparisons test, mCh: Pre vs Post, **p = 0.003. hM4Di: Pre vs Post, p = 0.1102. Right, direct comparison of CPP performance. Two-tailed, unpaired t-test, t_14_ = 2.04, p = 0.061. mCh, *n* = 8, hM4Di, *n* = 8.

**Extended Data Fig 6.**
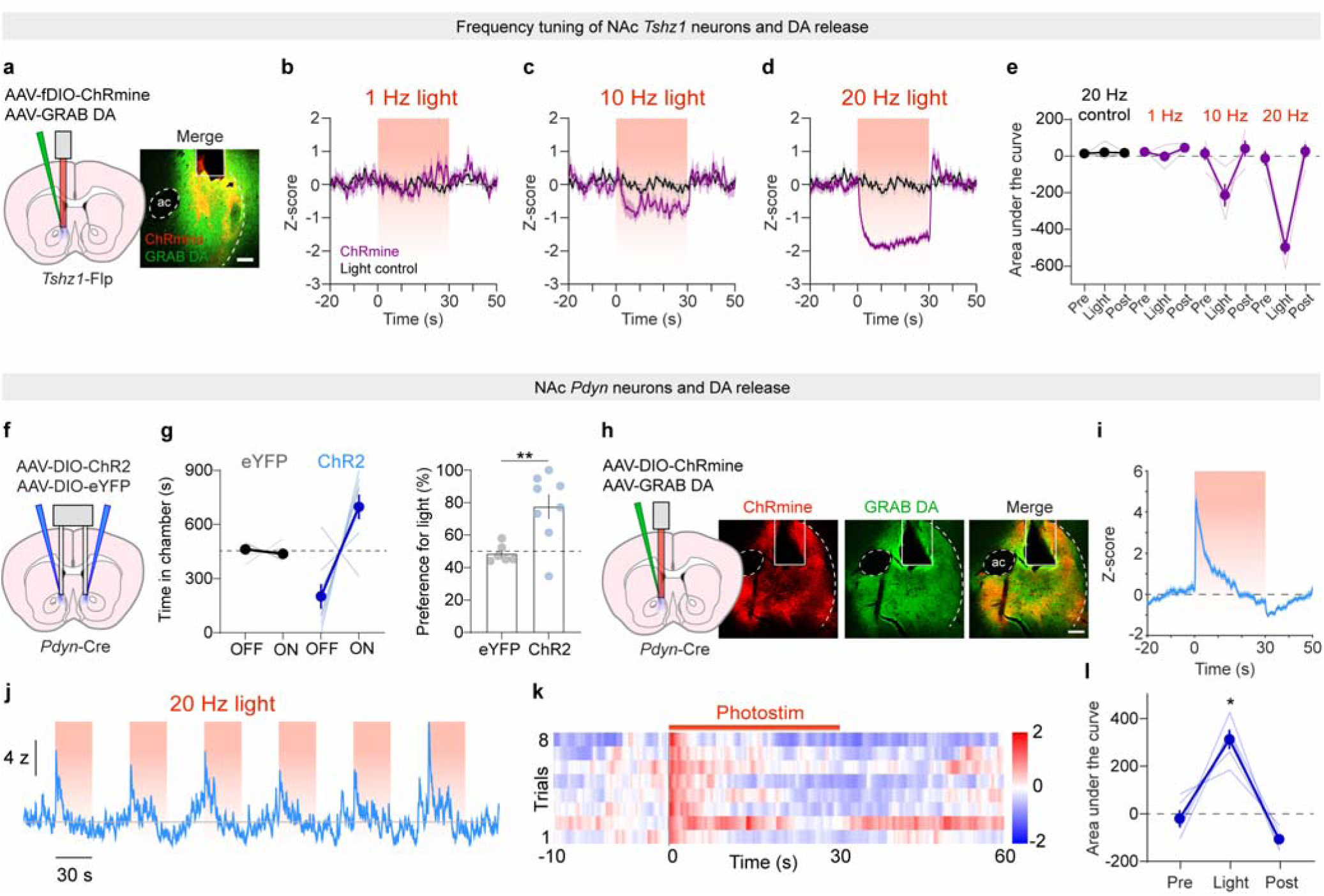
Frequency tuning of optogenetic stimulation of *Tshz1* neurons and stimulation of NAc *Pdyn* neurons. **a.** Photometry and optogenetic stimulation configuration. Scale = 100 um. **b.** Histogram showing the effect of 1 Hz stimulation of *Tshz1* neurons over a 30 second window, compared with a no opsin light control group. **c.** Histogram showing the effect of 10 Hz stimulation of *Tshz1* neurons over a 30 second window, compared with the same no opsin light control group. **d.** Histogram showing the effect of 20 Hz stimulation of *Tshz1* neurons over a 30 second window, compared with the same no opsin light control group. **e.** Area under the curve for each optogenetic treatment. **f.** Schematic of viral injections and fiber implants for real time place testing in *Pdyn*-Cre mice **g.** Left, time spent in the light-paired chamber for each group. RM two-way ANOVA, time x light, F_1,12_ = 10.25, **p = 0.0076. Sidak’s multiple comparisons test, eYFP: OFF vs ON, p = 0.9763. ChR2: OFF vs ON, **p = 0.0011. Right, direct comparison between groups. Unpaired, two-tailed t-test, t_12_ = 3.202, **p = 0.0076. eYFP, *n* = 6, ChR2, *n* = 8. **h.** Right, schematic of viral injection and fiber implant for simultaneous photometry recording and optical stimulation of NAc *Pdyn* neurons. Right, image of ChRmine and GRAB DA expression in the NAc. Scale = 100 um. **i.** Event histogram of average DA release while *Pdyn* neurons are stimulated with red light over a 30-second window. **j.** Representative trace across multiple *Pdyn* neuron activation windows demonstrating strong and consistent potentiation of local DA release. **k.** Color map showing the effect of photomstimulation of *Pdyn* neurons on DA release. **l.** Area under the curve quantification of the peri-event histogram presented in panel **i**. RM one-way ANOVA, F_2,12_ = 30.79, **p = 0.0036. Dunnett’s multiple comparison test, Pre vs Light, *p = 0.0187, Post vs Light, **p = 0.0036, Pre vs Post, *p = 0.0376. *n* = 5 mice.

**Extended Data Fig 7.**
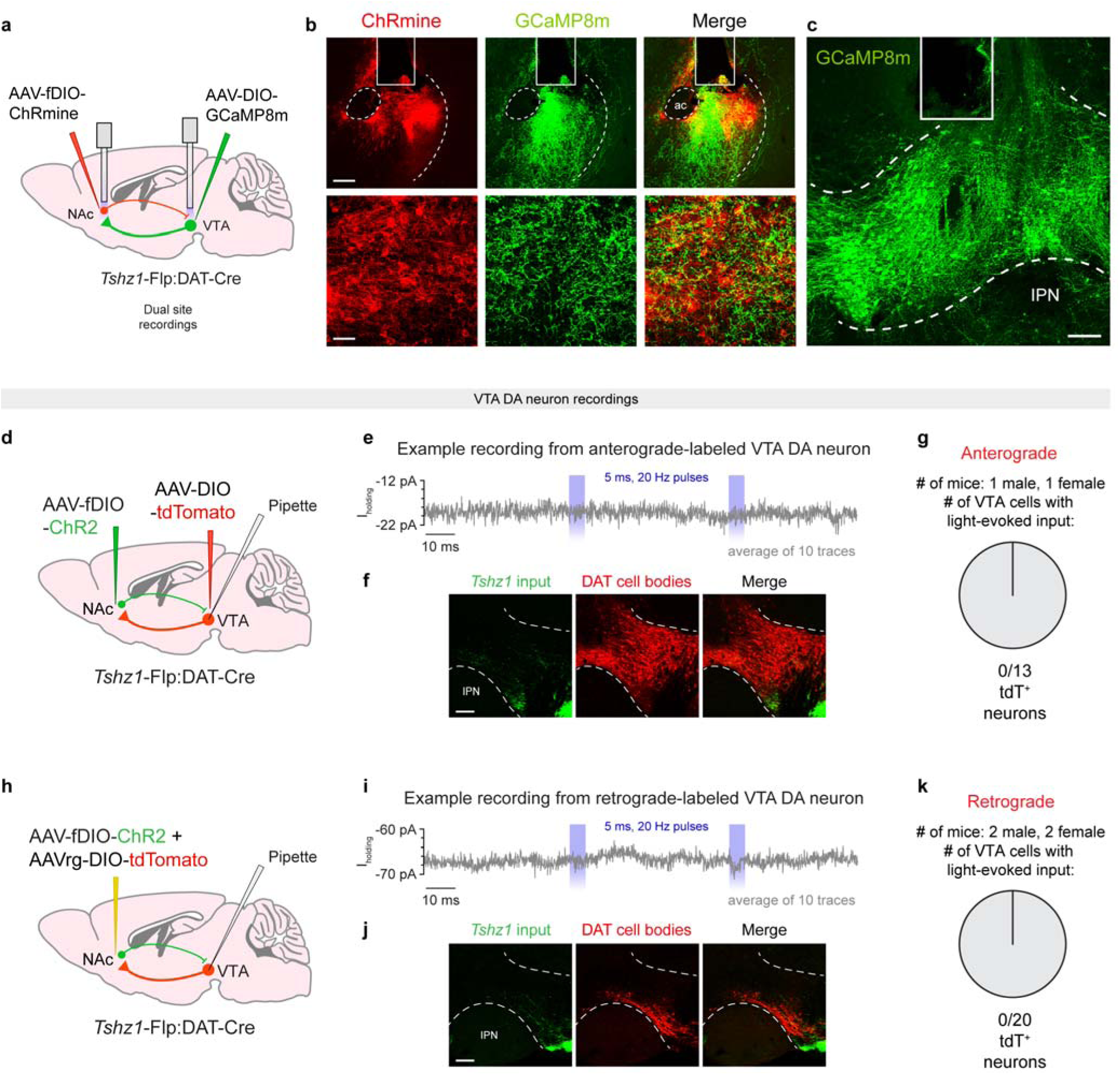
Dual photometry histology and whole-cell recordings of *Tshz1* inputs to VTA DA neurons. **a.** Dual recordings of VTA DA cell bodies and axons while NAc *Tshz1* cell bodies are stimulated. **b.** Representative images of injection and recording sites. Top left, image of NAc showing expression of ChRmine in *Tshz1* neurons and GCaMP8m in VTA DA axons. Scale = 100 um. Bottom left, high magnification image of ChRmine-infected *Tshz1* neurons surrounded by VTA DA axons. Scale = 50 um. **c.** Image of GCaMP8m expression in VTA DA cell bodies. Scale = 100 um. **d.** Viral injection strategy for anterograde-labeled VTA DA neurons. **e.** Example voltage clamp recording in a VTA DA neuron. Trace depicts average of ten sweeps from a single recording. Blue rectangles indicate time of two 5 ms LED pulses delivered 50 ms apart. **f.** Viral expression in the VTA using anterograde labeling strategy. **g.** Number of anterograde-labeled VTA DA neurons recorded (n = 13 tdTomato (tdT)^+^ neurons from 2 mice). **h.** Viral injection strategy for retrograde-labeled VTA DA neurons. **i.** Example voltage clamp recording in a VTA DA neuron. Trace depicts average of ten sweeps from a single recording. Blue rectangles indicate time of two 5 ms LED pulses delivered 50 ms apart. **j.** Viral expression in the VTA using retrograde labeling strategy. **k.** Number of retrograde-labeled VTA DA neurons recorded (n = 20 tdT^+^ neurons from 4 mice).

**Extended Data Fig 8.**
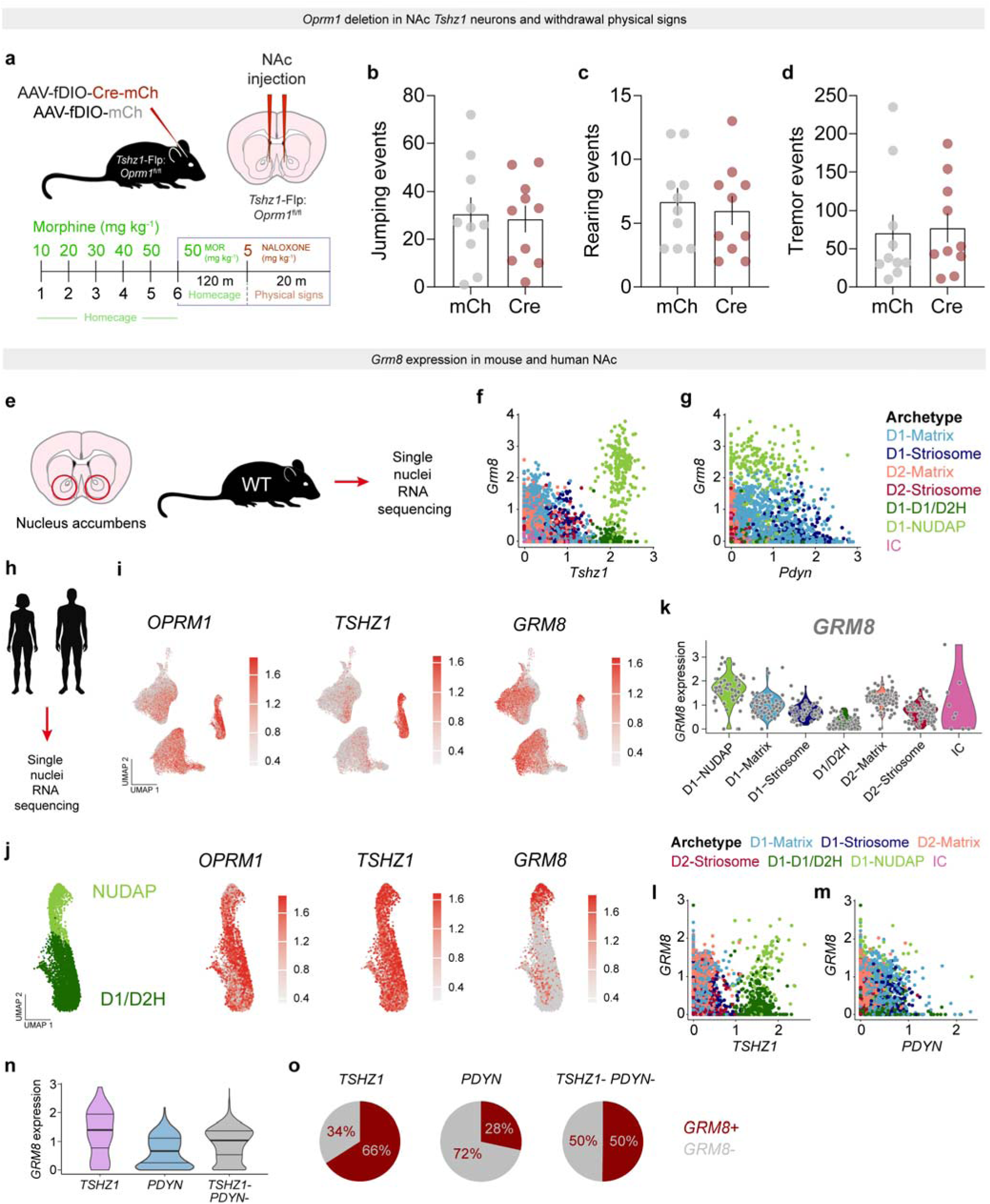
Impact of MOR KO in *Tshz1* neurons on withdrawal physical signs and *Grm8* expression analysis. **a.** Experimental design of withdrawal somatic sign testing. **b.** Quantification of jumping events during precipitated withdrawal. Two-tailed, unpaired t-test, t_18_ = 0.2355, p = 0.8165. **c.** Quantification of rearing events, two-tailed, unpaired t-test, t_18_ = 0.2126, p = 0.8340. **d.** Quantification of tremor events, two-tailed, unpaired t-test, t_18_ = 0.4476, p = 0.6598. mCh, *n* = 10, Cre, *n* = 10. **e.** snRNAseq in mouse NAc. **f.** Plot showing relationship between *Tshz1* and *Grm8* expression in all cells colored by archetype. D1-NUDAP cells stand out as containing high expression of both *Tshz1* and *Grm8*. **g.** Plot showing relationship between *Pdyn* and *Grm8* expression in all cells colored by archetype. **h.** snRNAseq in human NAc. **i.** Feature plots showing the expression pattern of *OPRM1*, *TSHZ1*, and *GRM8*. **j.** Expanded view of eccentric cluster feature plots showing the co-expression of *OPRM1*, *TSHZ1*, and *GRM8* in the NUDAP archetype. **k.** Violin plot quantifying the expression pattern of *GRM8* in each cell archetype. **l.** Plot showing relationship between *TSHZ1* and *GRM8* expression in all cells colored by archetype. D1-NUDAP cells stand out as containing high expression of both *TSHZ1* and *GRM8*. **m.** Plot showing relationship between *PDYN* and *GRM8* expression in all cells colored by archetype. **n.** Violin plot quantifying the expression pattern of *GRM8* in each cell-type. Mixed effect linear model, *TSHZ1* (NUDAP) vs *PDYN* ****p = 1.533e-15, *TSHZ1* (NUDAP) vs *TSHZ1*- *PDYN*-****p = 1.032e-90. *TSHZ1* (NUDAP) vs *PDYN,* Bonferroni corrected p = 3.066e-15. *TSHZ1* (NUDAP) vs *TSHZ1*- *PDYN*-, Bonferroni corrected p = 2.065e-90. **o.** Pie charts illustrating the proportion of each cell-type expressing *GRM8*.

## References

1 Blanco, C. & Volkow, N. D. Management of opioid use disorder in the USA: present status and future directions. Lancet 393, 1760–1772, doi:10.1016/S0140-6736(18)33078-2 (2019).

2 The Lancet Regional, H.-A. Opioid crisis: addiction, overprescription, and insufficient primary prevention. Lancet Reg Health Am 23, 100557, doi:10.1016/j.lana.2023.100557 (2023).

3 Volkow, N. D. & Blanco, C. The changing opioid crisis: development, challenges and opportunities. Mol Psychiatry 26, 218–233, doi:10.1038/s41380-020-0661-4 (2021).

4 Koob, G. F. A role for brain stress systems in addiction. Neuron 59, 11–34, doi:10.1016/j.neuron.2008.06.012 (2008).

5 Pantazis, C. B. et al. Cues conditioned to withdrawal and negative reinforcement: Neglected but key motivational elements driving opioid addiction. Sci Adv 7, doi:10.1126/sciadv.abf0364 (2021).

6 Darcq, E. & Kieffer, B. L. Opioid receptors: drivers to addiction? Nat Rev Neurosci 19, 499–514, doi:10.1038/s41583-018-0028-x (2018).

7 Fields, H. L. & Margolis, E. B. Understanding opioid reward. Trends Neurosci 38, 217–225, doi:10.1016/j.tins.2015.01.002 (2015).

8 Matthes, H. W. et al. Loss of morphine-induced analgesia, reward effect and withdrawal symptoms in mice lacking the mu-opioid-receptor gene. Nature 383, 819–823, doi:10.1038/383819a0 (1996).

9 Chaudun, F. et al. Distinct micro-opioid ensembles trigger positive and negative fentanyl reinforcement. Nature 630, 141–148, doi:10.1038/s41586-024-07440-x (2024).

10 Rossetti, Z. L., Hmaidan, Y. & Gessa, G. L. Marked inhibition of mesolimbic dopamine release: a common feature of ethanol, morphine, cocaine and amphetamine abstinence in rats. Eur J Pharmacol 221, 227–234, doi:10.1016/0014-2999(92)90706-a (1992).

11 Stella, L. et al. Interactive role of adenosine and dopamine in the opiate withdrawal syndrome. Naunyn Schmiedebergs Arch Pharmacol 368, 113–118, doi:10.1007/s00210-003-0773-9 (2003).

12 Gysling, K. & Wang, R. Y. Morphine-induced activation of A10 dopamine neurons in the rat. Brain Res 277, 119–127, doi:10.1016/0006-8993(83)90913-7 (1983).

13 Johnson, S. W. & North, R. A. Opioids excite dopamine neurons by hyperpolarization of local interneurons. J Neurosci 12, 483–488, doi:10.1523/JNEUROSCI.12-02-00483.1992 (1992).

14 Matsui, A. & Williams, J. T. Opioid-sensitive GABA inputs from rostromedial tegmental nucleus synapse onto midbrain dopamine neurons. J Neurosci 31, 17729–17735, doi:10.1523/JNEUROSCI.4570-11.2011 (2011).

15 Wise, R. A. & Bozarth, M. A. Action of drugs of abuse on brain reward systems: an update with specific attention to opiates. Pharmacol Biochem Behav 17, 239–243, doi:10.1016/0091-3057(82)90076-4 (1982).

16 Koob, G. F. Neurobiology of Opioid Addiction: Opponent Process, Hyperkatifeia, and Negative Reinforcement. Biol Psychiatry 87, 44–53, doi:10.1016/j.biopsych.2019.05.023 (2020).

17 Danjo, T., Yoshimi, K., Funabiki, K., Yawata, S. & Nakanishi, S. Aversive behavior induced by optogenetic inactivation of ventral tegmental area dopamine neurons is mediated by dopamine D2 receptors in the nucleus accumbens. Proc Natl Acad Sci U S A 111, 6455–6460, doi:10.1073/pnas.1404323111 (2014).

18 Steinberg, E. E. et al. Amygdala-Midbrain Connections Modulate Appetitive and Aversive Learning. Neuron 106, 1026–1043 e1029, doi:10.1016/j.neuron.2020.03.016 (2020).

19 Ungless, M. A., Magill, P. J. & Bolam, J. P. Uniform inhibition of dopamine neurons in the ventral tegmental area by aversive stimuli. Science 303, 2040–2042, doi:10.1126/science.1093360 (2004).

20 Zhu, Y., Wienecke, C. F., Nachtrab, G. & Chen, X. A thalamic input to the nucleus accumbens mediates opiate dependence. Nature 530, 219–222, doi:10.1038/nature16954 (2016).

21 Smith, A. C. W. et al. A master regulator of opioid reward in the ventral prefrontal cortex. Science 384, eadn0886, doi:10.1126/science.adn0886 (2024).

22 Aston-Jones, G., Hirata, H. & Akaoka, H. Local opiate withdrawal in locus coeruleus in vivo. Brain Res 765, 331–336, doi:10.1016/s0006-8993(97)00682-3 (1997).

23 Maldonado, R., Stinus, L., Gold, L. H. & Koob, G. F. Role of different brain structures in the expression of the physical morphine withdrawal syndrome. J Pharmacol Exp Ther 261, 669–677 (1992).

24 Rasmussen, K., Beitner-Johnson, D. B., Krystal, J. H., Aghajanian, G. K. & Nestler, E. J. Opiate withdrawal and the rat locus coeruleus: behavioral, electrophysiological, and biochemical correlates. J Neurosci 10, 2308–2317, doi:10.1523/JNEUROSCI.10-07-02308.1990 (1990).

25 Andraka, E., Phillips, R. A., 3rd, Brida, K. L. & Day, J. J. Chst9 marks a spatially and transcriptionally unique population of Oprm1-expressing neurons in the nucleus accumbens. Addict Neurosci 11, doi:10.1016/j.addicn.2024.100153 (2024).

26 Castro, D. C. & Bruchas, M. R. A Motivational and Neuropeptidergic Hub: Anatomical and Functional Diversity within the Nucleus Accumbens Shell. Neuron 102, 529–552, doi:10.1016/j.neuron.2019.03.003 (2019).

27 Frenois, F., Cador, M., Caille, S., Stinus, L. & Le Moine, C. Neural correlates of the motivational and somatic components of naloxone-precipitated morphine withdrawal. Eur J Neurosci 16, 1377–1389, doi:10.1046/j.1460-9568.2002.02187.x (2002).

28 Gracy, K. N., Dankiewicz, L. A. & Koob, G. F. Opiate withdrawal-induced fos immunoreactivity in the rat extended amygdala parallels the development of conditioned place aversion. Neuropsychopharmacology 24, 152–160, doi:10.1016/S0893-133X(00)00186-X (2001).

29 Stinus, L., Le Moal, M. & Koob, G. F. Nucleus accumbens and amygdala are possible substrates for the aversive stimulus effects of opiate withdrawal. Neuroscience 37, 767–773, doi:10.1016/0306-4522(90)90106-e (1990).

30 St Laurent, R., Martinez Damonte, V., Tsuda, A. C. & Kauer, J. A. Periaqueductal Gray and Rostromedial Tegmental Inhibitory Afferents to VTA Have Distinct Synaptic Plasticity and Opiate Sensitivity. Neuron 106, 624–636 e624, doi:10.1016/j.neuron.2020.02.029 (2020).

31 Gayden, J. et al. Integrative multi-dimensional characterization of striatal projection neuron heterogeneity in adult brain. bioRxiv, doi:10.1101/2023.05.04.539488 (2023).

32 Yao, Z. et al. A high-resolution transcriptomic and spatial atlas of cell types in the whole mouse brain. Nature 624, 317–332, doi:10.1038/s41586-023-06812-z (2023).

33 He, J. et al. Transcriptional and anatomical diversity of medium spiny neurons in the primate striatum. Curr Biol 31, 5473–5486 e5476, doi:10.1016/j.cub.2021.10.015 (2021).

34 Matsushima, A. et al. Transcriptional vulnerabilities of striatal neurons in human and rodent models of Huntington’s disease. Nat Commun 14, 282, doi:10.1038/s41467-022-35752-x (2023).

35 Fasano, L. et al. The gene teashirt is required for the development of Drosophila embryonic trunk segments and encodes a protein with widely spaced zinc finger motifs. Cell 64, 63–79, doi:10.1016/0092-8674(91)90209-h (1991).

36 Xiao, X. et al. A Genetically Defined Compartmentalized Striatal Direct Pathway for Negative Reinforcement. Cell 183, 211–227 e220, doi:10.1016/j.cell.2020.08.032 (2020).

37 Saunders, A. et al. Molecular Diversity and Specializations among the Cells of the Adult Mouse Brain. Cell 174, 1015–1030 e1016, doi:10.1016/j.cell.2018.07.028 (2018).

38 Bocklisch, C. et al. Cocaine disinhibits dopamine neurons by potentiation of GABA transmission in the ventral tegmental area. Science 341, 1521–1525, doi:10.1126/science.1237059 (2013).

39 Holly, E. N., Davatolhagh, M. F., Espana, R. A. & Fuccillo, M. V. Striatal low-threshold spiking interneurons locally gate dopamine. Curr Biol 31, 4139–4147 e4136, doi:10.1016/j.cub.2021.06.081 (2021).

40 Kramer, P. F. et al. Synaptic-like axo-axonal transmission from striatal cholinergic interneurons onto dopaminergic fibers. Neuron 110, 2949–2960 e2944, doi:10.1016/j.neuron.2022.07.011 (2022).

41 Kramer, P. F., Twedell, E. L., Shin, J. H., Zhang, R. & Khaliq, Z. M. Axonal mechanisms mediating gamma-aminobutyric acid receptor type A (GABA-A) inhibition of striatal dopamine release. Elife 9, doi:10.7554/eLife.55729 (2020).

42 Liu, C. et al. An action potential initiation mechanism in distal axons for the control of dopamine release. Science 375, 1378–1385, doi:10.1126/science.abn0532 (2022).

43 Mohebi, A. et al. Dissociable dopamine dynamics for learning and motivation. Nature 570, 65–70, doi:10.1038/s41586-019-1235-y (2019).

44 Gosnell, H. B. et al. mGluR8 modulates excitatory transmission in the bed nucleus of the stria terminalis in a stress-dependent manner. Neuropsychopharmacology 36, 1599–1607, doi:10.1038/npp.2011.40 (2011).

45 Schmid, S. & Fendt, M. Effects of the mGluR8 agonist (S)-3,4-DCPG in the lateral amygdala on acquisition/expression of fear-potentiated startle, synaptic transmission, and plasticity. Neuropharmacology 50, 154–164, doi:10.1016/j.neuropharm.2005.08.002 (2006).

46 Duvoisin, R. M. et al. Acute pharmacological modulation of mGluR8 reduces measures of anxiety. Behav Brain Res 212, 168–173, doi:10.1016/j.bbr.2010.04.006 (2010).

47 Palazzo, E., de Novellis, V., Rossi, F. & Maione, S. Supraspinal metabotropic glutamate receptor subtype 8: a switch to turn off pain. Amino Acids 46, 1441–1448, doi:10.1007/s00726-014-1703-5 (2014).

48 Strang, J. et al. Opioid use disorder. Nat Rev Dis Primers 6, 3, doi:10.1038/s41572-019-0137-5 (2020).

49 Kaoru, T. et al. Molecular characterization of the intercalated cell masses of the amygdala: implications for the relationship with the striatum. Neuroscience 166, 220–230, doi:10.1016/j.neuroscience.2009.12.004 (2010).

50 Kuerbitz, J. et al. Loss of Intercalated Cells (ITCs) in the Mouse Amygdala of Tshz1 Mutants Correlates with Fear, Depression, and Social Interaction Phenotypes. J Neurosci 38, 1160–1177, doi:10.1523/JNEUROSCI.1412-17.2017 (2018).

51 Totty, M. S. et al. Transcriptomic diversity of amygdalar subdivisions across humans and nonhuman primates. bioRxiv, doi:10.1101/2024.10.18.618721 (2024).

52 Asede, D., Doddapaneni, D. & Bolton, M. M. Amygdala Intercalated Cells: Gate Keepers and Conveyors of Internal State to the Circuits of Emotion. J Neurosci 42, 9098–9109, doi:10.1523/JNEUROSCI.1176-22.2022 (2022).

53 Hagihara, K. M. et al. Intercalated amygdala clusters orchestrate a switch in fear state. Nature 594, 403–407, doi:10.1038/s41586-021-03593-1 (2021).

54 Royer, S., Martina, M. & Pare, D. An inhibitory interface gates impulse traffic between the input and output stations of the amygdala. J Neurosci 19, 10575–10583, doi:10.1523/JNEUROSCI.19-23-10575.1999 (1999).

55 Wang, W. et al. Striatal mu-opioid receptor activation triggers direct-pathway GABAergic plasticity and induces negative affect. Cell Rep 42, 112089, doi:10.1016/j.celrep.2023.112089 (2023).

56 Pomrenze, M. B. et al. Modulation of 5-HT release by dynorphin mediates social deficits during opioid withdrawal. Neuron 110, 4125–4143 e4126, doi:10.1016/j.neuron.2022.09.024 (2022).

57 Tran, M. N. et al. Single-nucleus transcriptome analysis reveals cell-type-specific molecular signatures across reward circuitry in the human brain. Neuron 109, 3088–3103 e3085, doi:10.1016/j.neuron.2021.09.001 (2021).

58 Palucha-Poniewiera, A., Novak, K. & Pilc, A. Group III mGlu receptor agonist, ACPT-I, attenuates morphine-withdrawal symptoms after peripheral administration in mice. Prog Neuropsychopharmacol Biol Psychiatry 33, 1454–1457, doi:10.1016/j.pnpbp.2009.07.029 (2009).

59 Dong, J. et al. Patch and matrix striatonigral neurons differentially regulate locomotion. bioRxiv, doi:10.1101/2024.06.12.598675 (2024).

60 Lazaridis, I. et al. Striosomes control dopamine via dual pathways paralleling canonical basal ganglia circuits. Curr Biol 34, 5263–5283 e5268, doi:10.1016/j.cub.2024.09.070 (2024).

61 Al-Hasani, R. et al. Distinct Subpopulations of Nucleus Accumbens Dynorphin Neurons Drive Aversion and Reward. Neuron 87, 1063–1077, doi:10.1016/j.neuron.2015.08.019 (2015).

62 Koob, G. F. Neurobiological substrates for the dark side of compulsivity in addiction. Neuropharmacology 56 **Suppl 1**, 18–31, doi:10.1016/j.neuropharm.2008.07.043 (2009).

63 Koob, G. F. & Le Moal, M. Drug abuse: hedonic homeostatic dysregulation. Science 278, 52–58, doi:10.1126/science.278.5335.52 (1997).

64 Liu, J. & Schulteis, G. Brain reward deficits accompany naloxone-precipitated withdrawal from acute opioid dependence. Pharmacol Biochem Behav 79, 101–108, doi:10.1016/j.pbb.2004.06.006 (2004).

65 Eshel, N. et al. Striatal dopamine integrates cost, benefit, and motivation. Neuron 112, 500–514 e505, doi:10.1016/j.neuron.2023.10.038 (2024).

66 Siletti, K. et al. Transcriptomic diversity of cell types across the adult human brain. Science 382, eadd7046, doi:10.1126/science.add7046 (2023).

67 Hao, Y. et al. Dictionary learning for integrative, multimodal and scalable single-cell analysis. Nat Biotechnol 42, 293–304, doi:10.1038/s41587-023-01767-y (2024).

68 Bilous, M. et al. Metacells untangle large and complex single-cell transcriptome networks. BMC Bioinformatics 23, 336, doi:10.1186/s12859-022-04861-1 (2022).

69 Law, C. W., Chen, Y., Shi, W. & Smyth, G. K. voom: Precision weights unlock linear model analysis tools for RNA-seq read counts. Genome Biol 15, R29, doi:10.1186/gb-2014-15-2-r29 (2014).

70 You, Y. et al. Publisher Correction: Modeling group heteroscedasticity in single-cell RNA-seq pseudo-bulk data. Genome Biol 24, 112, doi:10.1186/s13059-023-02965-2 (2023).

71 Harding, S. D. et al. The IUPHAR/BPS Guide to PHARMACOLOGY in 2024. Nucleic Acids Res 52, D1438–D1449, doi:10.1093/nar/gkad944 (2024).

